# Lathosterol oxidase (sterol C5-desaturase) deletion confers resistance to amphotericin B and sensitivity to acidic stress in *Leishmania major*

**DOI:** 10.1101/2020.04.20.051540

**Authors:** Yu Ning, Cheryl Frankfater, Fong-Fu Hsu, Rodrigo P. Soares, Camila A. Cardoso, Paula M. Nogueira, Noelia Marina Lander, Roberto Docampo, Kai Zhang

## Abstract

Lathosterol oxidase (LSO) catalyzes the formation of C5-C6 double bond in the synthesis of various types of sterols in mammals, fungi, plants and protozoa. In *Leishmania* parasites, mutations in *LSO* or other sterol biosynthetic genes are associated with amphotericin B resistance. To investigate the biological roles of sterol C5-C6 desaturation, we generated a *LSO*-null mutant line (*lso*^*–*^) in *Leishmania major*, the causative agent for cutaneous leishmaniasis. *Lso*^*–*^ parasites lacked the ergostane-based sterols commonly found in wild type *L. major* and instead accumulated equivalent sterol species without the C5-C6 double bond. These mutant parasites were replicative in culture and displayed heightened resistance to amphotericin B. However, they survived poorly after reaching the maximal density and were highly vulnerable to the membrane-disrupting detergent Triton X-100. In addition, *lso*^*–*^ mutants showed defects in regulating intracellular pH and were hypersensitive to acidic conditions. They also had potential alteration in the carbohydrate composition of lipophosphoglycan, a membrane-bound virulence factor in *Leishmania*. All these defects in *lso*^*–*^ were corrected upon the restoration of LSO expression. Together, these findings suggest that the C5-C6 double bond is vital for the structure of sterol core, and while the loss of LSO can lead to amphotericin B resistance, it also makes *Leishmania* parasites vulnerable to biologically relevant stress.

**IMPORTANCE:** Sterols are essential membrane components in eukaryotes and sterol synthesis inhibitors can have potent effects against pathogenic fungi and trypanosomatids. Understanding the roles of sterols will facilitate the development of new drugs and counter drug resistance. Lathosterol oxidase (aka sterol C5-desaturase) is required for the formation of C5-C6 double bond in the sterol core structure in mammals, fungi, protozoans, plants and algae. Functions of this C5-C6 double bond are not well understood. In this study, we generated and characterized a lathosterol oxidase-null mutant in *Leishmania major*. Our data suggest that the C5-C6 double bond is vital for the structure and membrane-stabilizing functions of leishmanial sterols. In addition, our results imply that while mutations in lathosterol oxidase can confer resistance to amphotericin B, an important antifungal and antiprotozoal agent, the alteration in sterol structure leads to significant defects in stress response that could be exploited for drug development.

## INTRODUCTION

Leishmaniasis is the second most deadly parasitic disease after malaria with more than 12 million people infected worldwide (1). The causative agents belong to a group of trypanosomatid protozoans known as *Leishmania*. In the sandfly vector, *Leishmania* parasites are flagellated, extracellular promastigotes whereas in the mammalian host, they are non-flagellated, intracellular amastigotes (2). Current treatments are limited with toxic side effects and resistance is on the rise (3). Without a safe vaccine, it is necessary to identify new drug targets, develop new treatments, and decipher the mechanism of drug resistance in *Leishmania* (4).

The biosynthesis of sterol is an important pathway for most eukaryotes. In mammals, the dominant type of sterol is cholesterol, a vital membrane component which is also the precursor of steroid hormones (5). In fungi and trypanosomatids, ergostane-based sterols such as ergosterol and 5-dehydroepisterol are synthesized in high abundance which play equivalent roles as cholesterol in cellular membranes (6, 7). Ergosterol differs from cholesterol in the presence of two more double bonds: one at C7-C8 on the B ring and the other at C22-C23 on the side chain (Fig. S1)(8). In addition, ergosterol has an extra methyl group at the C24 position (Fig. S1). These structural differences make sterol biosynthesis a desirable source for antifungal and anti-trypanosomatid drug targets.

Amphotericin B (Amp B) is a polyene antibiotic that binds to ergostane-based sterols on the plasma membrane of pathogenic fungi or *Leishmania* leading to pore formation and accumulation of reactive oxygen species (ROS) (9-11). It has been used successfully to treat antimony-resistant leishmaniasis and in patients co-infected with *Leishmania* spp. and human immunodeficiency virus (11) (12). However, resistance to Amp B has been reported both in the laboratory and in clinical isolates (13-16). Multiple Amp B-resistant *Leishmania* lines show altered sterol composition and mutations in sterol biosynthetic enzymes such as the sterol C24-methyltransferase (SMT, EC 2.1.1.41) and sterol C14-alpha-demethylase (C14DM, EC 1.14.13.70) (13-16).

To interrogate the roles of these enzymes in *L. major*, we generated null mutants of C14DM (*c14dm*^*–*^) and SMT (*smt*^*–*^) using the targeted gene deletion approach (17). Both *c14dm*^*–*^ and *smt*^*–*^ mutants lack ergostane-based sterols but are viable in culture and highly resistant to Amp B (18, 19). *C14dm*^*–*^ mutants are extremely sensitive to heat and highly attenuated in virulence (19). They also display altered morphology, cytokinesis defects and increased plasma membrane fluidity (19). By comparison, defects exhibited by *smt*^*–*^ mutants, including elevated mitochondrial membrane potential and superoxide level, are less drastic (18). Interestingly, both *c14dm*^*–*^ and *smt*^*–*^ mutants show altered expression of lipophosphoglycan (LPG), a GPI-anchored virulence factor (20). Compared to *L. major* wild type (WT) parasites, the cellular level of LPG appears to be much lower in *c14dm*^*–*^ but higher in *smt*^*–*^ mutants (18, 19). These findings suggest that loss-of-function mutations in C14DM and SMT can lead to Amp B resistance but result in significant defects in stress response and virulence.

In addition to C14DM and SMT, mutations in the gene encoding lathosterol oxidase (LSO) are also implicated in Amp B resistance (15, 21, 22). LSO (aka sterol C5-desaturase) catalyzes the formation of C5-C6 double bond in the B ring of sterol intermediates, a late step in sterol synthesis (Fig. S1)(23). Orthologs of LSO have been identified in mammals, yeast, protozoans, plants and algae (15, 23-26). In *Saccharomyces cerevisiae*, the activity of LSO (encoded by *ERG3*) is sensitive to cyanide and requires iron, NAD(P)H as well as molecular oxygen (27). LSOs from yeast and *Tetrahymena thermophila* exbibit dependence on cytochrome b5 and cytochrome b5 reductase, suggesting that this desaturation reaction shares similarity to the sterol C4- and C14-demethylation steps (27-30) (Fig. S1).

In *S. cerevisiae*, deletion or inactivation of *ERG3/LSO* results in accumulation of episterol and depletion of ergosterol, and the null mutants fail to grow in the absence of heme synthesis (23, 31) (Fig. S1). In addition, *ERG3/LSO* mutants are unable to utilize respiratory substrates such as glycerol, acetate and ethanol (25). Furthermore, LSO expression contributes to tolerance to high temperature and acidic pH in fission yeast (32). These studies allude to the functions of LSO in regulating respiration and stress response in fungi although the mechanism of action is not well understood.

Studies on LSO in trypanosomatids are scarce. In light of its potential involvement in the development of Amp B resistance (15), it is necessary to characterize LSO in *Leishmania* and determine whether it is essential in the promastigote and amastigote stages, and whether it is required for the sensitivity of *Leishmania* to sterol synthesis inhibitors or Amp B. To address these questions, we generated LSO-null mutants in *L. major*. Our results indicate that LSO deletion confers resistance to Amp B but the mutants show poor survival and reduced growth rate under the acidic condition due to their inability to maintain intracellular pH. In addition, LSO deletion altered the structure of LPG and reduced parasite virulence in mice. These findings shed new light on the roles of sterol synthesis in *Leishmania* stress response and reveal the fitness costs associated with the development of drug resistance.

## RESULTS

### Genetic deletion and cellular localization of *L. major* LSO

The putative *L. major LSO* gene (TriTrypDB: LmjF.23.1300) is located on chromosome 23, showing 40% identity to *S. cerevisiae* ERG3p (Gene ID: 850745) and 38% identity to human sterol C5-desaturase (GenBank: BAA33729.1). Based on the predicted open reading frame (ORF), *L. major* LSO protein contains 302 amino acids with four transmembrane helices, no obvious signal peptide and is expected to catalyze the formation of a double bond between C5 and C6 in the B ring of sterol intermediates (Fig. S1).

To investigate the roles of sterol C5-C6 desaturation reaction in *L. major*, we replaced the endogenous *LSO* alleles with nourseothricin (*SAT*) and blasticidin (*BSD*) resistance genes using the homologous recombination approach (17). The resulting LSO-null mutants (*lso*^*–*^) were verified by Southern blot with an ORF probe and a 5’-flanking sequence probe (Fig. 1A). To complement the null mutants, we introduced a LSO-expressing plasmid (pXG-LSO) into *lso*^*–*^ to generate *lso*^*–*^*/+LSO* (the add-back strain). To examine the cellular localization of LSO, the C-terminus of LSO was fused to GFP and introduced into *lso*^*–*^ (*lso*^*–*^*/+LSO-GFP*) (Fig. S2A). In immunofluorescence microscopy, LSO-GFP showed a similar distribution (∼72% overlap) as BiP, an endoplasmic reticulum (ER) marker (33) (Fig. 1B-F), suggesting that LSO is primarily located at the ER. This result is similar to the localizations of C14DM and SMT in *Leishmania* (18, 19), as well as LSO in *T. thermophila* (28).

**Figure 1.**
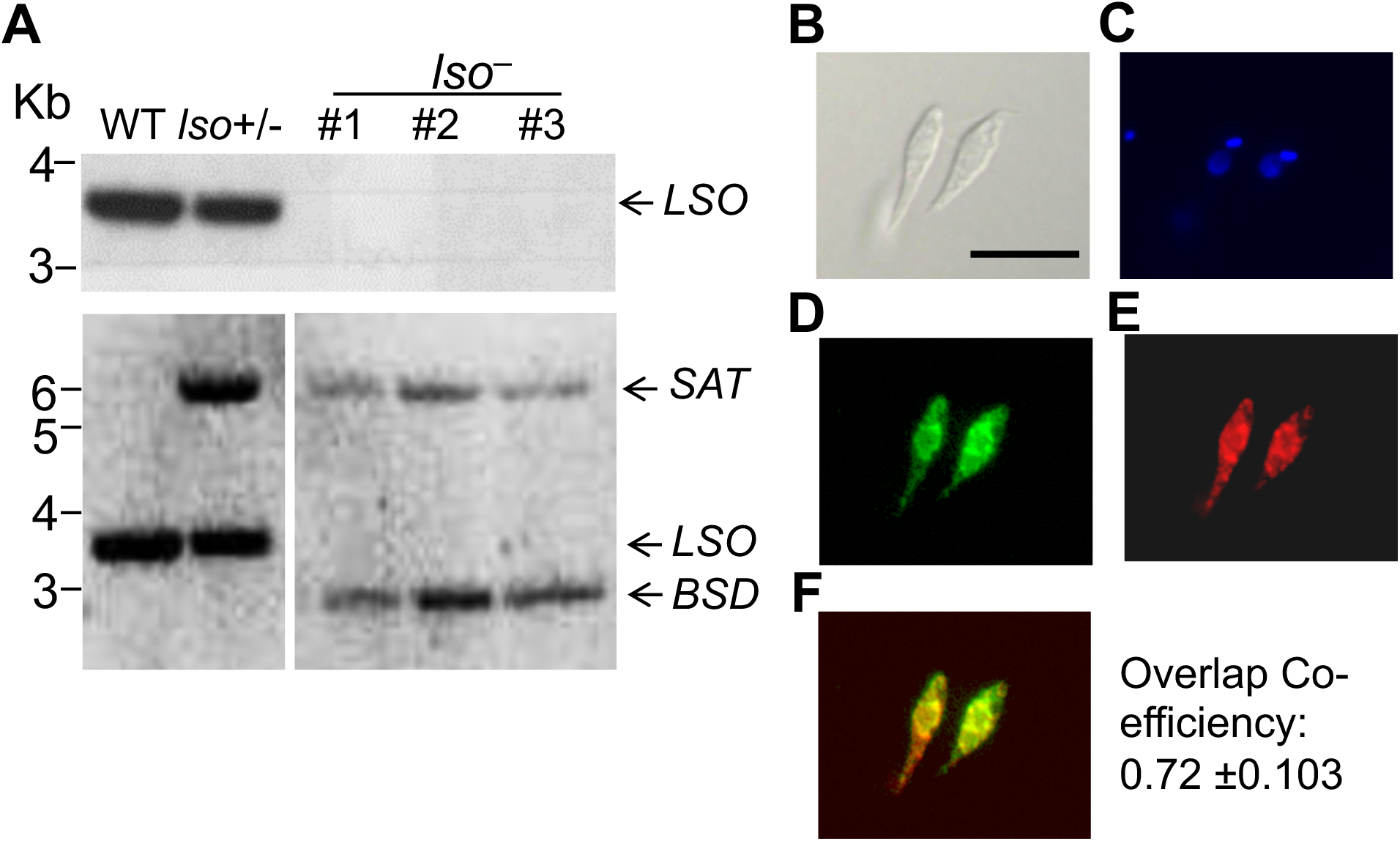
Genetic knockout and cellular localization of LSO. (**A**) Genomic DNA samples from *L. major* LV39WT, *LSO* +/-(heterozygous knockout), and *lso*^*–*^ (homozygous knockout clones 1-3) parasites were processed for Southern blot analyses, using radioactive probes from the open reading frame (top) or upstream flanking region of *LSO* (bottom). Bands corresponding to *LSO* and drug resistance genes (*BSD/SAT*) are indicated. (**B**-**F**) Immunofluorescence microscopy of *lso*^*–*^/+LSO-GFP promastigotes. (**B**) Differential interference contrast (Scale bar: 10 µm); (**C**) DNA staining with Hoechst; (**D**) GFP fluorescence; (**E**) Anti-BiP (an ER marker) staining followed by goat anti-rabbit IgG-Texas Red; and (**F**) Merge of **D** and **E**. Overlap between GFP and ER was calculated using JaCOP Image J analysis from 30 cells.

### *Lso*^*–*^ promastigotes have altered sterol composition

Sterols from promastigotes were converted into trimethylsilyl (TMS) derivatives, followed by electron ionization (EI) gas chromatography-mass spectrometry (GC-MS) analysis. Sterol molecules were identified based on their molecular weights, retention times, and EI spectra. Consistent with the findings we previously reported, *L. major* WT promastigotes contained two main sterols, i.e. ergosterol and 5-dehydroepisterol, represented by peaks 1 and 2 in Fig. 2A, respectively (18, 19). Interestingly, all the sterols from *lso*^*–*^ promastigotes were shifted to the right in the GC chromatogram including two dominant peaks 1’ and 2’ (Fig. 2B and Fig. S2B-C). While ergosterol and 5-dehydroepisterol had the expected molecular weight of 468.5 as TMS derivatives, the dominant sterols from *lso*^*–*^ (1’ and 2’) had the molecular weight of 470.5 as TMS derivatives (Fig. 2D). By library search, peak 1’ and peak 2’ matched ergosta-7,22-dien-3-ol, (3β,22E)- and episterol-TMS derivatives, respectively. While we did not have the ergosta-7,22-dien-3-ol, (3β,22E) standard to confirm the structure of peak 1’, the retention time and the EI mass spectrum of peak 2’ in *lso*^*–*^ were identical to those of the episterol standard (Fig. 2D-F). These findings are consistent with role of LSO in catalyzing the C5-C6 desaturation reaction to form 5-dehydroepisterol (Fig. S1). GC-MS analysis on the area ratios of internal standard peak and leishmanial sterol peaks revealed no significant difference in total cellular sterol abundance between WT and *lso*^*–*^promastigotes. Importantly, introduction of LSO or LSO-GFP into *lso*^*–*^ restored the sterol profile to WT-like composition (Fig. 2C and Fig. S2). Taken together, these results support the identity of *L. major* LSO as a sterol C5-desaturase.

**Figure 2.**
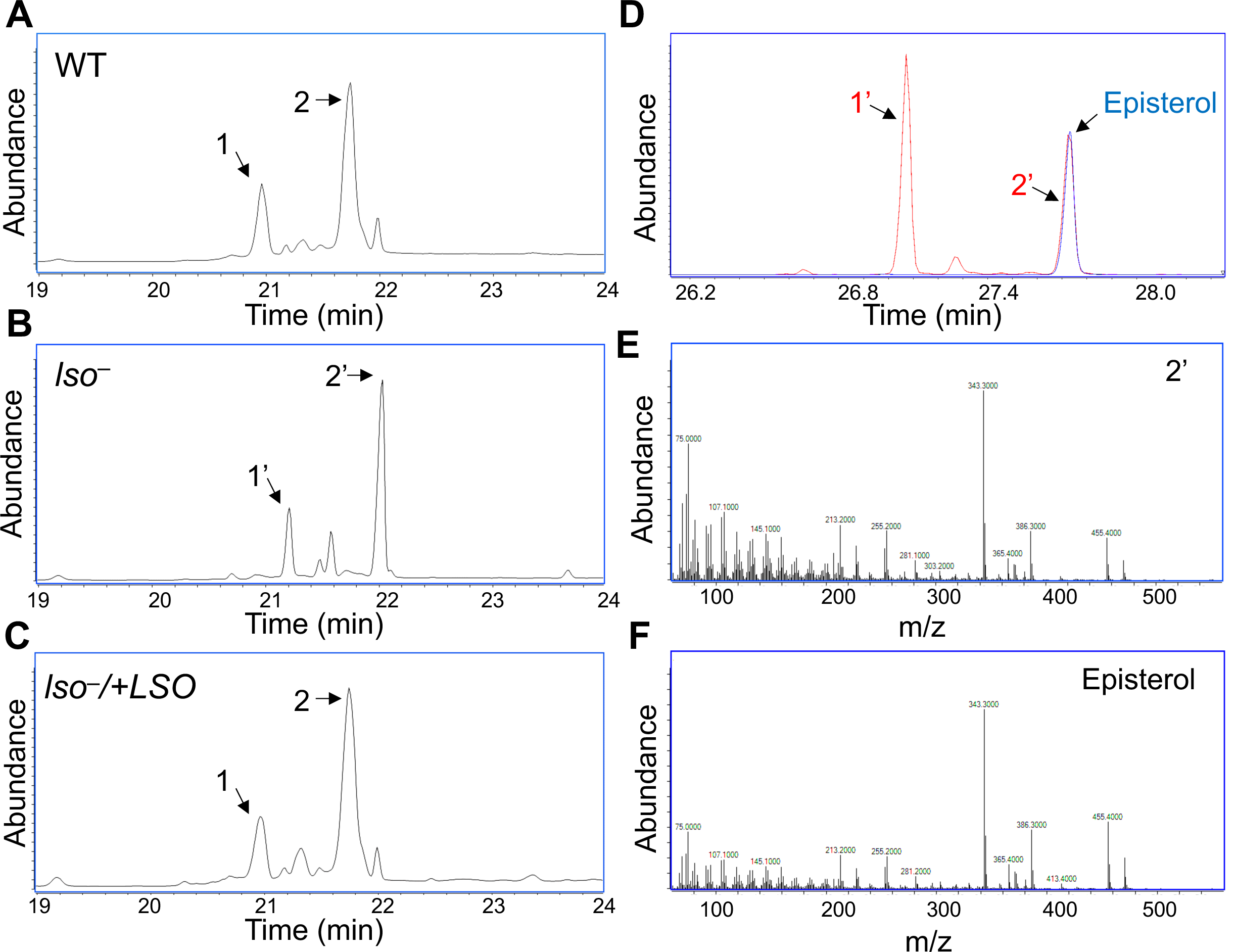
GC-MS analyses of sterol TMS derivatives show altered sterol profile in *lso*^*–*^ mutants. (**A-C**) Total ion current chromatograms plotted from 19.0-24.0 min scans (mass range: m/z 50-550) of the sterol TMS derivatives of (**A**) LV39WT; (**B**) *lso*^*–*^; and (**C**) *lso*^*–*^/*+LSO*. In **A** and **C**, peaks 1 and 2 represent ergosterol and 5-dehydroepisterol, respectively. In **B**, the peaks including 1’ and 2’ are shifted to the right. (**D**) The reconstructed ion chromatogram of the M^+^ ion (m/z 470.5) from full GC-MS scans (m/z 50-550) of the *lso*^*–*^ sample (trace in red) and from the episterol standard (trace in blue). In addition to the perfect match of the retention time of peak 2’ with the episterol standard, the full scan EI mass spectra (70 eV) plotted from peak 2’ (**E**) and the episterol standard (**F**) are also identical, confirming that peak 2’ is episterol.

### *Lso*^*–*^ mutants are replicative in culture but show poor survival in stationary phase

The *LSO*-null mutants were fully viable in culture with a doubling time of ∼8 hours during the log phase and could reach a maximal density of 2-3 × 10^7^ cells/ml, similar to WT and add-back parasites (Fig. 3A). However, after reaching maximal density, *lso*^*–*^ promastigotes showed significantly reduced viability in the stationary phase. First, we measured the percentage of cells whose long axis was less than twice the length of the short axis. Such a “round” shape was indicative of cells under duress. In early stationary phase (stationary day 1-2), 18-32% of *lso*^*–*^ promastigotes were round whereas only 2-9% of WT and add-back cells were round (Fig. 3B). The difference became less pronounced in late stationary stage (stationary day 3-4) when the percentages of round cells increased among WT and add-back parasites (Fig. 3B). Similarly, we observed a much higher percentage of dead cells in *lso*^*–*^ mutants (28-38%) than WT and add-back parasites (3-14%) from stationary day 2 to day 3 (Fig. 3C). We also examined the ability of *lso*^*–*^ mutants to form metacyclics, which are the non-dividing and infective form of promastigotes (34). *Lso*^*–*^ produced 40-50% less metacyclics than WT and add-back parasites in stationary phase. In conclusion, LSO is not required for the survival or replication of log phase promastigotes but is important for maintaining viability during the stationary phase.

**Figure 3.**
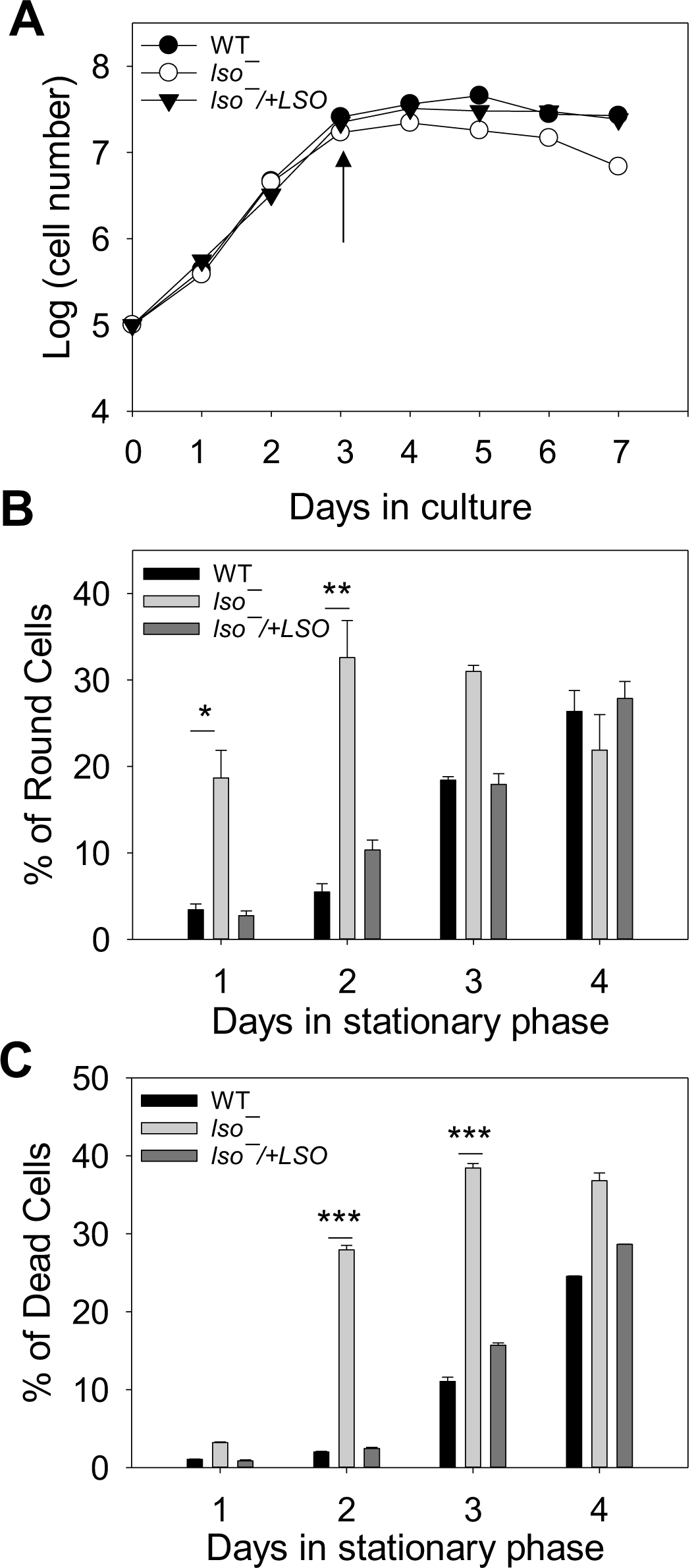
*Lso*^*–*^ mutants show poor survival in stationary phase. (**A**) Promastigotes were inoculated in M199 media at 1 x 10^5^/ml and cell densities were determined daily. The arrow marks the onset of stationary phase. In (**B**)-(**C**), percentage of round cells (**B**) and dead cells (**C**) were measured during day 1-4 in stationary phage. Error bars represent standard deviations from 3 experiments (**: *p* < 0.01, ***: *p* < 0.001).

### *Lso*^*–*^ promastigotes show increased resistance to Amp B and are hypersensitive to Triton X-100

Amp B is a potent drug that interacts with ergosterol or ergosterol-like sterols on the plasma membrane resulting in pore formation, oxidant accumulation and cell death (9-11, 35). Alterations in sterol biosynthesis can confer resistance to Amp B (14, 16). In *Candida lusitaniae*, the Amp B-resistant clinical isolates showed reduced *ERG3* gene expression, suggesting that the C5-C6 double bond contributes to the binding of Amp B to membrane sterol (22). In another report, mutations in sterol C5-desaturase (LSO) were found to be associated with Amp B resistance in *Leishmania mexicana* (15). Here we measured the sensitivity of *lso*^*–*^ promastigotes to Amp B in liquid culture by growing cells in various concentrations of Amp B for 48 hours (Fig. 4A). The effective concentrations to inhibit 25%, 50% and 90% of growth (EC25, EC50 and EC90) were determined using cells grown in the absence of Amp B as a control (Table 1). In comparison to WT and *lso*^*–*^*/+LSO* promastigotes, *lso*^*–*^ mutants were 2-4 times more resistant to Amp B (Fig. 4A and Table 1). The increase in Amp B resistance was close to the *smt*^*–*^ mutants (18) but not as pronounced as the *c14dm*^*–*^ mutants (10-100 times more resistant than WT)(19). This result suggests that C5-C6 desaturation enhances the binding between membrane sterol and Amp B. Meanwhile, the susceptibility of *lso*^*–*^ mutants to itraconazole, an inhibitor of C14DM (19), was similar to WT and *lso*^*–*^*/+LSO* parasites (Fig. 4B).

**Table 1.** *Lso*^*–*^ mutants show reduced sensitivity to Amp B.

| EC25 (μM±SD) |  |  | EC50 (μM±SD) |  |  | EC90 (μM±SD) |  |  |
| --- | --- | --- | --- | --- | --- | --- | --- | --- |
| WT | *lso*<sup>-</sup> | WT/*lso*<sup>-</sup> | WT | *lso*<sup>-</sup> | WT/*lso*<sup>-</sup> | WT | *lso*<sup>-</sup> | WT/*lso*<sup>-</sup> |
| 0.034±<br>0.003 | 0.077±<br>0.004 | **1/2.24** | 0.046±<br>0.003 | 0.133±<br>0.005 | **1/2.89** | 0.084±<br>0.005 | 0.303±<br>0.033 | **1/3.61** |

EC25, EC50 and EC90 (Average ± standard deviations from 3 independent experiments) were calculated through comparison to control cells grown in absence of Amp B.

**Figure 4.**
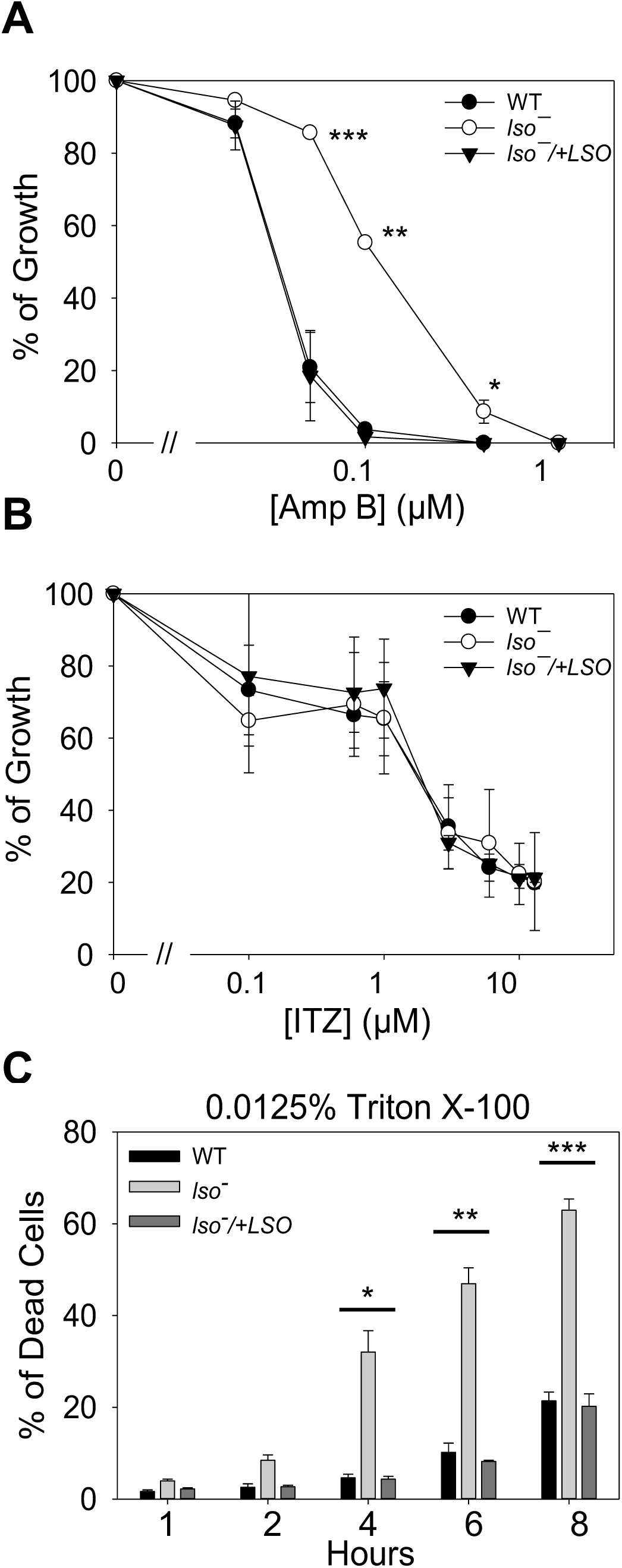
*Lso*^*–*^ mutants are resistant to Amp B and hypersensitive to Triton X-100. (**A-B**) Log phase promastigotes were inoculated into M199 media with different concentrations of Amp B (**A**) or ITZ (**B**). Cells grown in absence of drugs were used as controls and percentages of growth were calculated after 48 hours. (**C**) Log phase promastigotes were incubated in the presence of 0.0125% Triton X-100 for 1-8 hours. Cell viability was determined by flow cytometry. Error bars represent standard deviations from 3-4 experiments (*: *p* <0.05, **: *p* <0.01, ***: *p* <0.001).

We also examined whether the change in sterol composition could alter the plasma membrane stability in *lso*^*–*^ mutants. Log phase promastigotes were incubated in regular media containing 0.0125% of Triton X-100 and percentages of dead cells were monitored over time. As shown in Fig. 4C, *lso*^*–*^ exhibited hypersensitivity to Triton X-100 after 4 hours and the defect was rescued by the introduction of *LSO*. This finding resembles our previous observation in the *c14dm*^*–*^ mutants which are unable to form detergent-resistant membrane fractions (19). Therefore, alteration in sterol composition may cause hypersensitivity to detergent-induced plasma membrane disruption in *L. major*.

### LSO is required for promastigote survival and optimal growth under acidic condition

When promastigotes are transmitted from the sandfly vector to the mammalian host, they encounter elevated temperature, acidic pH and oxidative bursts (36). Here we investigated whether LSO is required for parasites to survive under stress conditions. First, promastigotes were inoculated in complete media at pH 7.4 (the regular pH) or pH 5.0 for 0-60 hours to examine their tolerance to acidic stress. While WT and *lso*^*–*^*/+LSO* cells showed good viability at pH 5.0 (< 8% death), 25-30% of *lso*^*–*^ mutants died after 8-12 hours (Fig. 5A-B). The dead cell percentage in *lso*^*–*^ went down after 24 hours (although still higher than WT and *lso*^*–*^*/+LSO*) which was likely due to the rapid lysis of dead cells (Fig. 5B). Next, we evaluated the ability of *lso*^*–*^ promastigotes to withstand heat stress by increasing the culture temperature from 27 to 37. As indicated in Fig. 5C, no significant difference was detected until 48 hours into the temperature shift when *lso*^*–*^ mutants showed ∼ 2 times more dead cells than WT and add back parasites. In addition, we also incubated promastigotes in phosphate buffered saline (PBS) to assess their resistance to starvation, and results indicated that the *lso*^*–*^ mutants responded similarly to WT parasites (Fig. 5D).

**Figure 5.**
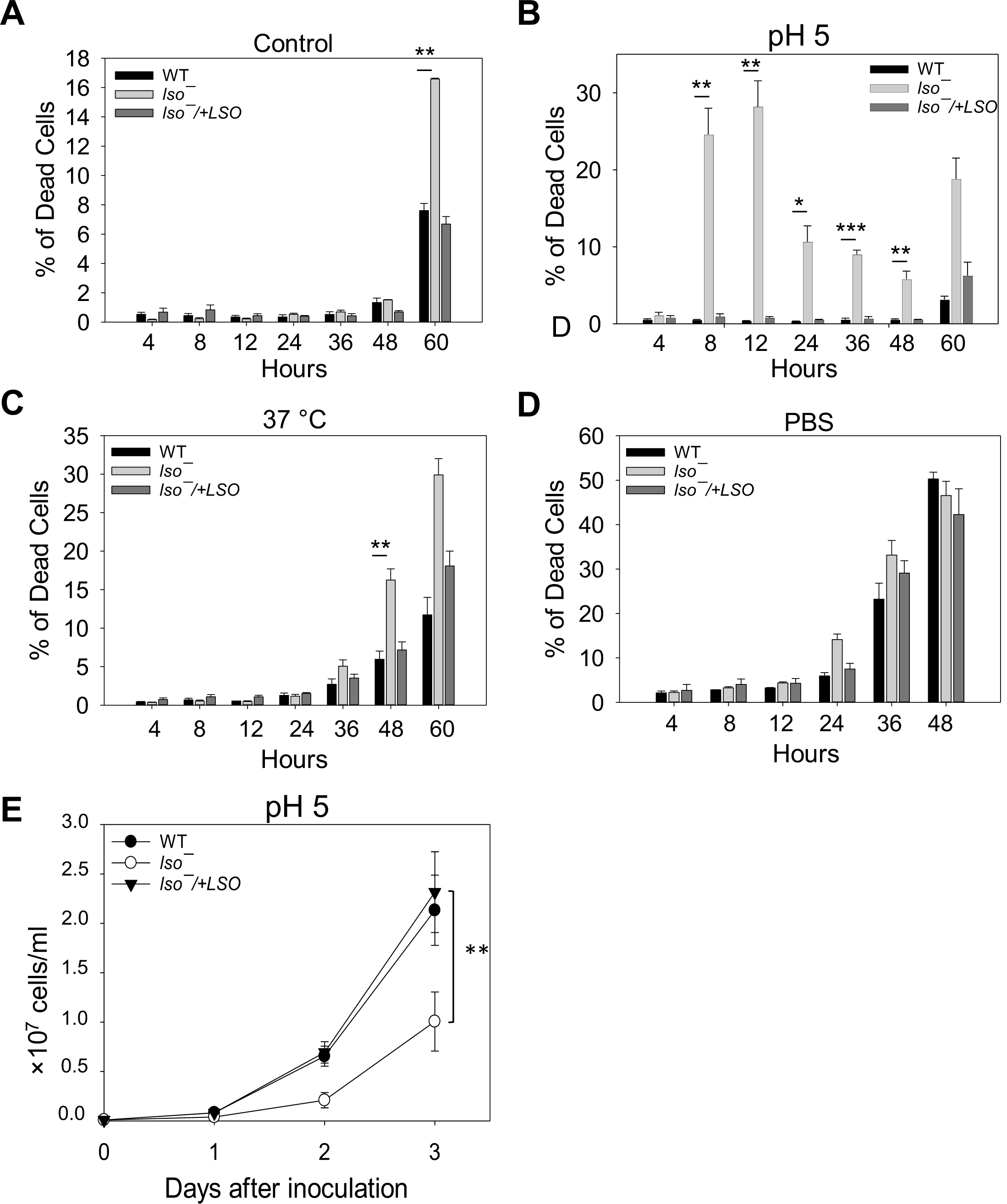
*Lso*^*–*^ mutants show poor viability under acidic and heat stress. (**A**)-(**D**) Log phase promastigotes were incubated at 5 x 10^6^ cells/ml in M199 media (**A-C**) or PBS (**D)** under neutral (**A, C** and **D**) or acidic conditions (**B**) at either 27 □(**A, B**, and **D**) or 37 □ (**C**). Percentages of dead cells were determined at the indicated times. (**E**) Log phase promastigotes were inoculated into an acidic medium (pH 5.0) at 1 x 10^5^ cells/ml and culture densities were determined daily. Error bars represent standard deviations from 4 experiments (*: *p* <0.05, **: *p* <0.01, ***: *p* <0.001).

Consistent with their hypersensitivity to acidic pH, *lso*^*–*^ promastigotes proliferated slowly in the pH 5.0 medium (Fig. 5E). Overall, the mutants’ poor survival and growth delay at pH 5 suggest that LSO is crucial for *L. major* promastigotes to tolerate acidic stress and may affect parasites growth in the phagolysosome (37).

### *LSO* contributes to intracellular pH homeostasis

Their hypersensitivity to acidic condition prompted us to examine whether *lso*^*–*^ mutants can regulate intracellular pH. When cultivated in the regular medium (pH 7.4), *lso*^*–*^ had a slightly lower intracellular pH than WT parasites (7.5 vs 7.8) (Table 2). However, when grown in an acidic medium (pH 5.0), the intracellular pH of *lso*^*–*^ dropped to 7.0 whereas in WT and *lso*^*–*^ */+LSO* parasites it remained at 7.8-7.9 (Table 2). This finding argues that alteration in sterol composition can affect the cytosolic pH homeostasis in *L. major*.

**Table 2.** LSO is required for the maintenance of intracellular pH.

| Cell type | pH 7.4 |  | pH 5.0 |  |
| --- | --- | --- | --- | --- |
|  | Mean | SD | Mean | SD |
| WT | 7.76 | 0.265 | 7.90 | 0.122 |
| *lso*<sup>-</sup> | 7.48 | 0.265 | 7.05 | 0.031 |
| *lso*<sup>-</sup>/+*LSO* | 7.83 | 0.124 | 7.78 | 0.130 |

Promastigotes were cultivated in the regular medium (pH 7.4) or acidic medium (pH 5.0) for 3 days and intracellular pH values were measured after labeling with 10 μM of BCECF-AM for 30 min. Mean values and standard deviations were calculated from 3 experiments (\*\*\*: $p < 0.001$ ).

Plasma membrane is the first barrier against the change of extracellular pH. Hypersensitivity of *lso*^*–*^ mutants to Triton X-100 (Fig. 4C) suggests that their plasma membrane is less stable, thus affecting their ability to control intracellular pH. Besides plasma membrane, certain intracellular organelles and proteins are essential for intracellular pH homeostasis as well (38). Acidocalcisomes are electron-dense acidic organelles rich in divalent cations and polyphosphate first identified in trypanosomatids (39). They play important roles in calcium homeostasis, osmoregulation and the maintenance of intracellular pH (39, 40). The acidocalcisome has a vacuolar-type H^+^-pyrophosphatase (VP1) which is involved in the uptake of H^+^ from the cytosol into acidocalcisomes and regulates intracellular pH homeostasis (41, 42). After labeling stationary phase or metacyclic-like promastigotes with an anti-*T. brucei* VP1 antibody, we observed a 40-45% reduction in fluorescence intensity in *lso*^*–*^ mutants in comparison to WT and add-back parasites (Fig. 6A-B), suggesting that LSO is involved in the expression and/or localization of VP1 at the acidocalcisome. We did not detect any significant difference in acidocalcisome morphology, abundance or contents in short chain and long chain polyphosphate between *lso*^*–*^ and WT parasites (Fig. S3A-B and data not shown). From these analyses, we postulate that compromised expression of VP1 along with increased plasma membrane instability in *lso*^*–*^ may contribute to their inability to maintain intracellular pH when challenged with acidic stress.

**Figure 6.**
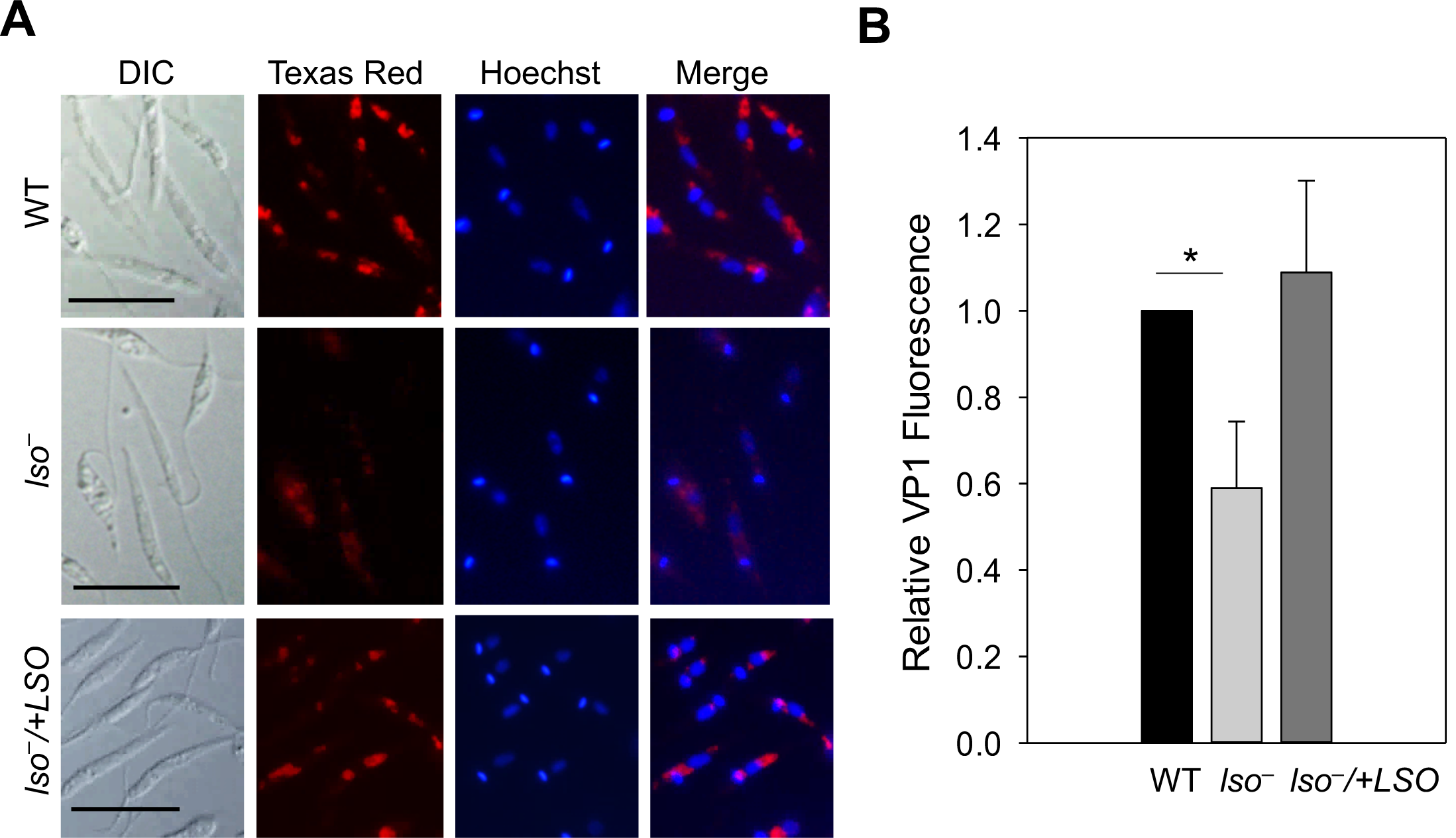
*Lso*^*–*^ mutants express less vacuolar proton pyrophosphatase (VP1). (**A**) Day 3 stationary phase promastigotes were labeled with rabbit anti-*Tb*VP1 antiserum (1:800), followed by anti-rabbit IgG-Texas Red (1:1000). DNA was stained with Hoechst. DIC: Differential interference contrast; Merge: the merge of Texas Red and Hoechst. Scale bar: 10 µm. (**B**) Relative levels of VP1 staining were determined from 100 metacyclic-like promastigotes for each line. Error bars represent standard deviations from 3 experiments (*: *p* <0.05).

### Depletion of *LSO* alters the expression and structure of lipophosphoglycan (LPG)

Sterols are enriched in the ordered microdomains (lipid rafts) along with sphingolipids and glycosylphosphatidylinositol (GPI)-anchored proteins on the plasma membrane (43). Previous work on *c14dm*^*–*^ and *smt*^*–*^ mutants demonstrates that alteration of sterol synthesis can affect the expression level of membrane-bound GPI-anchored virulence factors such as LPG and GP63 (a metalloprotease) (18, 19). To determine the role of LSO in the synthesis of GPI-anchored glycoconjugates, we performed western blot and immunofluorescence microscopy assays using the WIC79.3 monoclonal antibody which recognizes the terminal Gal (β1,3) subunits on the side chains branching off the Gal (β1,4)-Man (α1)-PO_4_ repeat units of *L. major* LPG (44, 45). As illustrated in Fig. 7A-B, whole cell lysates from log phase and stationary phase of *lso*^*–*^ appeared to have less LPG compared to WT and *lso*^*–*^*/+LSO* parasites (10-20% of WT-level, Fig. 7E). The reduction was not due to increased release of LPG from *lso*^*–*^ into the culture supernatant (Fig. 7A-B). Immunofluorescence microscopy assay confirmed this finding in *lso*^*–*^ mutants (Fig. 7G-H). Using the same WIC79.3 antibody, WT and add-back parasites displayed robust surface labeling at the exposure time of 100 ms (Fig. 7G). In contrast, signals from *lso*^*–*^ mutants were only detectable at a longer exposure time showing significant intracellular staining (Fig. 7H). Therefore, the expression of LPG was clearly altered in *lso*^*–*^. Meanwhile, these mutants showed similar level of GP63 as WT and add-back parasites (Fig. 7C-D and F).

**Figure 7.**
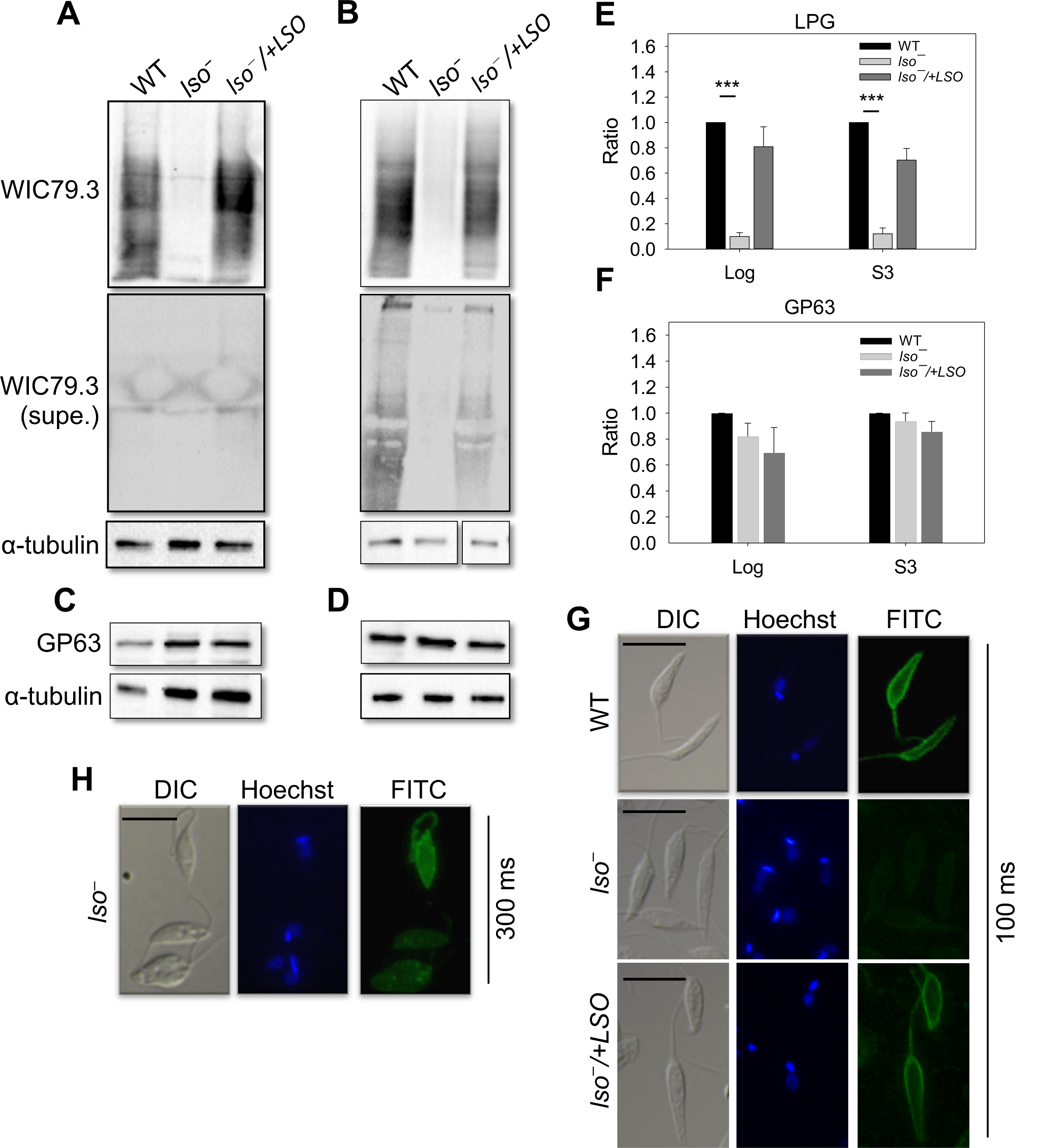
LSO is required for the synthesis of WT-like LPG. (**A-D**) Whole cell lysates or culture supernatants (supe.) from log phase (**A, C**) or day 3 stationary phase (**B, D**) promastigotes were analyzed by Western blot using anti*-L. major* LPG (mAb WIC79.3), anti-GP63 or anti-α-tubulin antibodies. (**E-F**) The relative abundance of LPG (**E**) and GP63 (**F**) in *lso*^*−*^ and *lso*^*−*^/*+LSO* were normalized to the levels in LV39WT promastigotes. Error bars represent standard deviations from 4 experiments (***: *p* <0.001). (**G**-**H**) Log phase promastigotes were labeled with mAb WIC79.3, followed by anti-mouse IgG-FITC. DNA was stained with Hoechst. Exposure times for the FITC channel: 100 ms for **G** and 300 ms for **H**. Scale bar: 10 µm.

LPG in *Leishmania* is a complex, polymorphic molecule composed of a lysoalkylphosphatidylinositol lipid anchor, a phosphorylated oligosaccharide core, a phosphoglycan backbone made of Gal (β1,4)-Man (α1)-PO_4_ repeat units which may contain side chains, and an oligosaccharide cap (20, 46). The lack of reactivity to WIC79.3 antibody in *lso*^*–*^ could reflect a loss or modification of side chains that branch off the phosphoglycan backbone (47, 48), or deficiencies in the synthesis of lipid anchor, oligosaccharide core or phosphoglycan backbone (49, 50). To probe the LPG structure in *lso*^*–*^, we first carried out a western blot analysis using the CA7AE monoclonal antibody, which recognizes the unsubstituted (bare) Gal (β1,4)-Man (α1)-PO_4_ backbone (51). As shown in Fig. 8A, CA7AE could label the LPG and related proteophosphoglycan from *L. donovani* (strains lS2D and LV82) which was expected since their phosphoglycan backbones were devoid of side chains (52). Meanwhile, no significant labeling was detected from *L. major* WT or *lso*^*–*^ parasites with the CA7AE antibody, suggesting that their LPG backbones had side chain modifications (Fig. 8A).

**Figure 8.**
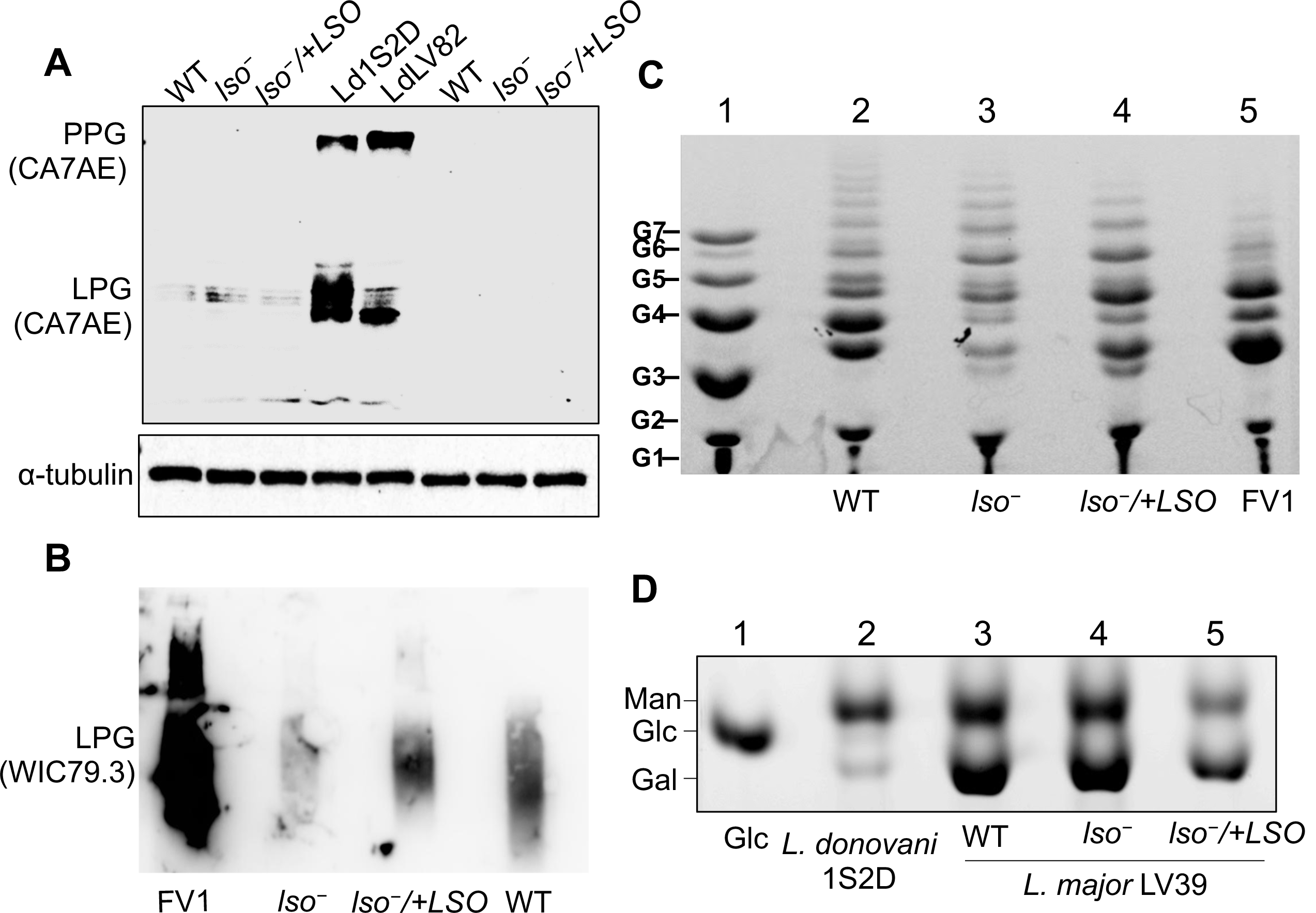
*Lso*^*–*^ mutants have less short side chain sugar residues on their LPG. (**A**) Whole cell lysates were processed for Western blot with mAb CA7AE or anti-α-tubulin antibody. Log phase (left three lanes) and day 3 stationary phase (right three lanes) promastigotes of LV39WT, *lso*^*−*^ and *lso*^*−*^/*+LSO* were analyzed along with *L. donovani* strain 1S2D and LV82 (middle two lanes). PPG: proteophosphoglycan. (**B**) Immunoblotting of purified LPG (5 µg per lane) from *L. major* strains probed with mAb WIC79.3. (**C**) FACE analysis of dephosphorylated LPG repeat units from *L. major* (lane 2-5). Lane 1: malto-oligomer ladder represented by 1–7 glucose residues (G1–G7). (**D**) Monosaccharide profile of GIPLs. Lane 1: glucose standard; lane 2: type I GIPL *from L. donovani* 1S2D containing mostly mannose residues and low levels of galactose; lane 3: repeat units of *L. major* LV39 WT (type II GIPL); lane 4: repeat units of *L. major lso*^*−*^; and lane 5: repeat units of *L. major lso*^*−*^/*+LSO*. Man: mannose; Glc: glucose; Gal: galactose. Experiments were performed twice and results from one representative set were shown.

To explore the carbohydrate composition of LPG side chain from *lso*^*–*^ mutants, we purified LPG from WT, *lso*^*–*^, and *lso*^*–*^/+*LSO* promastigotes as previously described (53). The yield of LPG was similar (150-200 μg/10^10^ cells for all lines) and these LPG samples exhibited similar reactivity to the WIC79.3 antibody as we observed with whole cell lysates (Fig. 7A-B and Fig. 8B). As expected, LPG from the *L. major* Friedlin V1 strain (positive control) was recognized strongly by this antibody (Fig. 8B) (48).

Next the LPG samples were subjected to fluorophore-assisted carbohydrate electrophoresis (FACE) to analyze the sizes of their Gal (β1,4)-Man (α1)-PO_4_ repeat units. As shown in Fig. 8C, *L. major* LV39 WT parasites had both short (G3-G4) and intermediate (G5-G11) side chains branching-off the LPG repeat units, indicative of a mixture of mono-, di- and poly-galactosylated residues on the side chains (47). Notably, *lso*^*–*^ mutants had a similar profile as LV39 WT for the intermediate side chains (G5-G11), but their short side chains (G3-G4) were much reduced (Fig. 8C). As expected, *L. major* FV1 parasites had more short side chains (G3-G4) than intermediate side chains (G5-G11) which is consistent with the dominance of Gal1-2 short side chains capped with arabinose in this strain (Fig. 8C) (48, 54, 55).

Next, we examined if LSO deletion affected the carbohydrate composition of glycoinositolphospholipids (GIPLs), another major glycoconjugate in *Leishmania* (56). After strong acid hydrolysis, the carbohydrate composition of GIPLs was very similar among WT, *lso*^*–*^, and *lso*^*–*^/+*LSO* promastigotes consistent with the galactose- and mannose-rich type II GIPLs (56, 57) (Fig. 8D). This profile was distinct from the type I GIPLs in *L. donovani* that is highly enriched in mannose (57) (Fig. 8D). Together these data suggest that LSO deletion does not change the carbohydrate profile of GIPLs but reduces the abundance of short (Gal1-2) side chains on the LPG backbone in *L. major*.

### *Lso*^*–*^ mutants have minor defects in mitochondrial functions

Previous reports indicate that inhibition of sterol biosynthesis can lead to compromised mitochondrial functions in trypanosomatids (18, 58-60). In *S. cerevisiae*, LSO/Erg3 is not required for viability in media containing ergosterol but mutants fail to grow on non-fermentable substrates such as glycerol and ethanol, suggesting that this enzyme is needed for respiration (31, 61). To assess the role of LSO in mitochondrial functions in *L. major*, we first examined the mitochondrial membrane potential (ΔΨ_m_) after labeling cells with tetramethylrhodamine ethyl ester (TMRE) (62). Compared to WT and add-back parasites, *lso*^*–*^ mutants had 30-50% higher ΔΨ_m_ in the early stationary stage but did not show significant difference in log phase or late log phase (Fig. S4A). To measure the production of mitochondrial ROS, we labeled cells using a mitochondria-specific ROS indicator MitoSox Red. As shown in Fig. S4B, *lso*^*–*^ mutants had slightly higher fluorescence signal than the WT and *lso*^*–*^*/+LSO* parasites (the difference was not statistically significant except for log phase), suggesting a modest accumulation of ROS in their mitochondria. Next, we used the MitoXpress probe to examine oxygen consumption rate (63) by incubating log phase promastigotes in a respiration buffer containing sodium pyruvate but no glucose. Under this condition, *lso*^*–*^ showed a slightly lower oxygen consumption rate than WT and add-back parasites although the difference was not statistically significant (Fig. S4C). Together, these data indicate that LSO deletion has minor effects on the mitochondrial functions in *L. major*.

### *Lso*^*–*^ mutants show attenuated virulence in a mouse model

To study the role of LSO in *L. major* virulence, metacyclics were isolated from stationary phase cultures and used to infect BALB/c mice in the footpads. Parasite virulence was assessed by measuring the development of footpad lesions over time. Compared to WT and *lso*^*–*^*/+LSO* parasites, mice infected by *lso*^*–*^ mutants showed a 2-4 weeks delay in lesion progression (Fig. 9A), which was consistent with the lower parasite numbers in the infected footpads at weeks 6 and 14 post infection (Fig. 9B). To explore the virulence of amastigotes, we isolated amastigotes from promastigote-infected footpads and used them to infect naive BALB/c mice. As shown in Fig. 9C-D, *lso*^*–*^ amastigotes were slightly attenuated in virulence in comparison to WT and *lso*^*–*^ */+LSO* amastigotes but the difference was less pronounced than metacyclics. These findings suggest that LSO is important for *L. major* promastigotes to grow and cause disease in mice.

**Figure 9.**
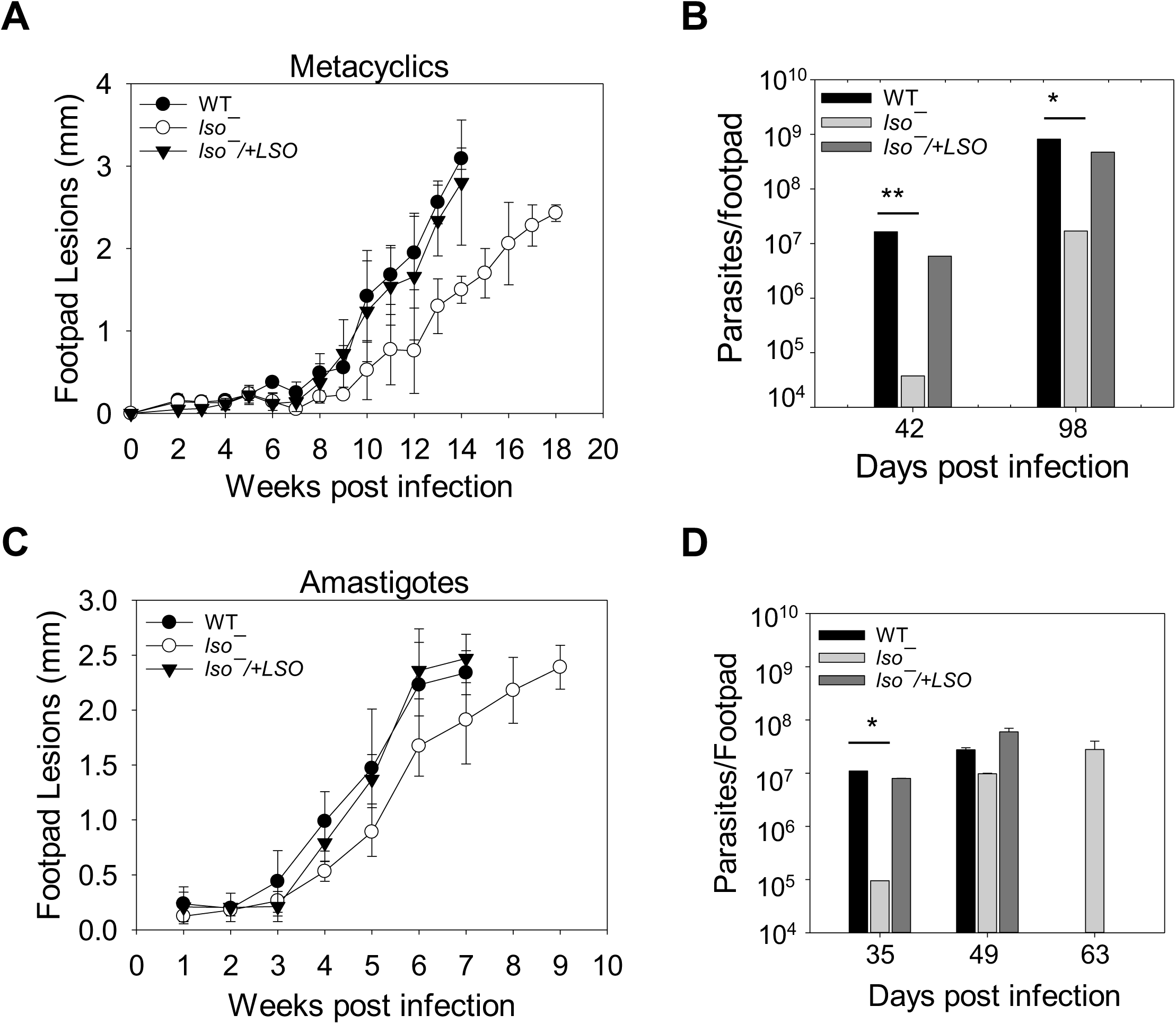
*Lso*^*–*^ mutants show attenuated virulence in mice. BALB/c mice were infected in the footpads with metacyclics (**A-B**) or amastigotes (**C-D**). Footpad lesions were measured weekly (**A** and **C**) and parasite numbers were determined by the limiting dilution assay (**B** and **D**). Error bars represent standard deviations (*: *p* <0.05, **: *p* <0.01). **TABLES**

## DISCUSSION

In this study, we characterized the gene encoding LSO (sterol C5-desaturase), a sterol biosynthetic enzyme, in the protozoan parasite *L. major*. LSO catalyzes the formation of a double bond between C5 and C6 in the B ring of sterol intermediates (Fig. S1). *L. major* LSO-null mutants (*lso*^*–*^) were devoid of ergosterol or 5-dehydroepisterol (abundant in WT parasites). Instead they accumulated C5-C6 saturated sterols such as ergosta-7,22-dienol and episterol (Fig. 2 and Fig. S2B-E). While the difference appears to be minor in chemical composition, *lso*^*–*^ mutants were 2-4 times more resistant to Amp B than WT and *lso*^*–*^*/+LSO* (Fig. 4A and Table 1). These mutants were fully replicative in culture during the log phase with minor mitochondrial defects but showed poor viability after entering stationary phase (Fig. 3 and Fig. S4). LSO deletion also led to hypersensitivity to acidic pH (Fig. 5). These observations are largely in agreement with the phenotypes of *ERG3* mutant in *S. cerevisiae* (23, 31).

It is interesting that LSO activity appears to contribute to the binding affinity between Amp B and ergostane-based sterols. The C5-C6 double bond is conserved among all major sterols including cholesterol (mammals), ergosterol (fungi and trypanosomatids) and stigmasterol (plants) (15, 23-26). Without this double bond, the A ring could twist/rotate more freely from the B ring, potentially making the sterol core less flat and reducing its binding capacity to Amp B (Fig. S1). Such a change in the sterol core conformation could also increase the gap between sterol and phospholipid, making the membrane less stable. This is consistent with the increased sensitivity of *lso*^*–*^ mutants to heat and Triton X-100 (Fig. 4C and Fig. 5C).

Compared to WT and add-back parasites, *lso*^*–*^ mutants had a lower intracellular pH and the difference became more pronounced when cells were cultivated in a pH 5.0 medium (Table 2). This finding is consistent with their reduced capacity to survive and replicate under the acidic condition (Fig. 5B and 5E). Besides affecting the plasma membrane, the loss of C5-C6 double bond in bulk sterol may influence the function of intracellular organelles as sterols are located not only in the plasma membrane but also in the membrane of intracellular organelles. Acidocalcisomes are membrane-enclosed storage organelles involved in osmoregulation, phosphate metabolism, calcium homeostasis, and intracellular pH maintenance in protozoan parasites (39). The expression of VP1, an acidocalcisome-associated vacuolar H^+^-pyrophosphatase, was lower in *lso*^*–*^ during the stationary phase (Fig. 6). Since VP1 transports protons from the cytosol into acidocalcisomes (using pyrophosphate hydrolysis as the energy source), the reduced VP1 expression may lead to a more acidic intracellular pH and slower recovery of intracellular pH under acidic conditions (42).

Similar to the C14DM-null mutants, the cellular level of LPG in *lso*^*–*^ appeared to be significantly reduced based on western blot and immunofluorescence microscopy analyses using the WIC79.3 antibody which recognizes the terminal Gal (β1,3) subunits on the side chain of *L. major* LPG (44, 45) (Fig. 7 and Fig. 8B). However, further analyses suggest that *lso*^*–*^ mutants still synthesized bulk glycoconjugates as their LPG and GIPLs could be extracted and purified to similar yields as WT and add-back parasites (LPG: ∼150 μg/10^10^ cells; GIPLS: ∼60 μg/10^10^ cells). Lack of reactivity to CA7AE antibody indicates that the LPG backbone in *lso*^*–*^ is not bare like in *L. donovani* (52)(Fig. 8A). In addition, FACE result indicates that *lso*^*–*^ mutants seemed to possess less short side chains and similar level of intermediate side chains compared to WT parasites (Fig. 8C). Together, these data imply that the low WIC79.3 reactivity in *lso*^*–*^ is not due to a total loss of LPG synthesis like several previously characterized LPG-synthetic mutants (49, 50, 64). Instead, it is likely caused by the reduced level of terminal Gal (β1,3) subunits on the LPG side chain. In *L. major*, certain groups of galactosyltransferases and arabinosyltransferases catalyze the attachment of galactose and arabinose to the LPG side chains, respectively (47, 48). Future studies on the expression of these sugar transferases, along with detailed LPG structure determination in sterol mutants (*lso*^*–*^, *c14dm*^*–*^ and *smt*^*–*^) will help elucidate the molecular mechanism by which sterol synthesis influences LPG production in *Leishmania*.

*Lso*^*–*^ promastigotes and amastigotes showed slightly reduced virulence in BALB/c mice (Fig. 9). The hypersensitivity to acidic pH and heat likely compromised their ability to survive and replicate in the phagolysosome (37). Alteration in LPG structure may also contribute to the virulence attenuation (65). Overall, the fitness loss displayed by *lso*^*–*^ is similar (in severity) to *smt*^*–*^ and less dramatic than *c14dm*^*–*^ mutants (18, 19). Nonetheless, modifications in sterol structure, even relatively minor changes (like in *lso*^*–*^), can significantly reduce parasite’s fitness.

In summary, our study shed new light into the biological roles of sterol C5-C6 desaturation in *Leishmania* growth, stress response and virulence. Along with previous reports on *smt*^*–*^ and *c14dm*^*–*^ mutants (18, 19), these findings reveal the potential fitness costs associated with mutations conferring Amp B resistance and may offer possible strategies to counter the development of drug resistance.

## MATERIALS AND METHODS

### Materials

M199 medium, cholesta-3,5-diene, phenyl-Sepharose CL-4B, and alkaline phosphatase (from *Escherichia coli*) were purchased from Sigma-Aldrich (St. Louis, MO). Itraconazole (ITZ) and amphotericin B (Amp B) were purchased from LKT Laboratories, Inc. (St. Paul, MN) and EMD Chemicals, Inc. (San Diego, CA), respectively. MitoXpress oxygen probe was purchased from Luxcel Biosciences (Cork, Ireland). AG50W-X12 cation-exchange and AG1-X8 anion-exchange resins were purchased from Bio-Rad (Hercules, CA). All other chemicals were purchased from Thermo Fisher Scientific unless specified otherwise.

### Molecular constructs

The predicted open reading frame (ORF) of *L. major LSO* (LmjF.23.1300, 302 aa) was amplified by PCR from *L. major* LV39 WT genomic DNA with primer #649 and #650 (Table S1). The PCR product was digested with BamHI and ligated into the pXG vector to generate pXG-*LSO*. A modified ORF of *LSO* was amplified by using primer #649 and #651 to remove the stop codon and then used to generate pXG-*LSO*-GFP for localization study.

To generate knockout constructs, the upstream and downstream flanking sequences (∼550 bp each) of *LSO* were amplified with primer pairs #645/#646 and #647/#659, respectively. These flanking sequences were digested and ligated into the cloning vector pUC18. Genes conferring resistance to nourseothricin (*SAT*) and blasticidin (*BSD*) were inserted between the upstream and downstream flanking sequences to generate pUC18-KO-*LSO::SAT* and pUC18-KO-*LSO::BSD*. All the molecular constructs were confirmed by restriction enzyme digestion and/or sequencing. Oligonucleotides used in this study are summarized in Table S1.

### *Leishmania* culture and genetic manipulation

Unlike otherwise specified, *L. major* strain LV39 clone 5 (Rho/SU/59/P), *L. major* strain Friedlin V1 (MHOM/IL/80/Friedlin), *L. donovani* strain 1S2D (MHOM/SD/00/1S-2D), and *L. donovani* strain LV82 (MHOM/ET/67:LV82) promastigotes were cultivated at 27 °C in M199 medium (pH 7.4, with 10% fetal bovine serum and other supplements) (66). The infective metacyclic promastigotes (metacyclics) were isolated from day 3 stationary phase promastigotes using the density centrifugation method (67). To generate the *LSO*-null mutants, the two *LSO* alleles in *L. major* LV39 WT parasites were replaced with *BSD* and *SAT* by homologous recombination (17). The resulting heterozygous (*LSO +/-*) and homozygous (*lso*^*–*^) mutants were confirmed by Southern blot. Briefly, genomic DNA samples were digested and resolved on a 0.7% agarose gel, transferred to a nitrocellulose membrane and hybridized with [^32^P]-labeled DNA probes targeting the ORF or a 500 bp upstream region of endogenous *LSO*. To restore *LSO* expression, pXG-*LSO* or pXG-*LSO*-GFP were introduced to *lso*^*–*^ by stable transfection, resulting in *lso*^*–*^/+*LSO* or *lso*^*–*^/+*LSO*-*GFP* parasites respectively.

### Sterol analysis by gas chromatography-mass spectrophotometry (GC-MS)

Total lipids were extracted from mid log phase promastigotes (3-7 x 10^6^ cells/ml) following the Folch’s method (68). An internal standard, cholesta-3,5-diene (FW: 368.34), was provided at 2.0 × 10^7^ molecules/cell during extraction. Lipid samples were dissolved in methanol at 1.0 × 10^9^ cells equivalence/ml. Equal amounts of each lipid extracts in methanol were transferred to separate vial inserts, evaporated to dryness under nitrogen and derivatized with 50 μl of BSTFA + 1% TMCS:acetonitrile (1:3) followed by heating at 70 °C for 30 min. GC-MS analysis was conducted on an Agilent 7890A GC coupled with Agilent 5975C MSD in electron ionization mode. Derivatized samples (2 μl each) were injected with a 10:1 split into the GC column with the injector and transfer line temperatures set at 250 °C. The GC temperature started at 180 °C, held for 2 min, followed by 10 °C/min increase until 300 °C, and held for 15 min. To confirm that the unknow GC peak retention time matched that of the episterol standard, we also used a second temperature program started at 80 °C for 2 min, ramped to 260 °C at 50 °C/min, held for 15 min, and increased to 300 °C at 10 °C/min and held for 10 min. A 25 m Agilent J & W capillary column (DB-1, 0.25 mm id, 0.1 μm film thickness) was used for the separation.

### Immunofluorescence microscopy

For LSO-GFP localization, *lso*^*–*^/+*LSO*-GFP parasites were labeled with rabbit anti-*T. brucei* BiP antiserum (1:2000) for 20 min followed by goat anti-rabbit IgG-Texas Red antibody (1:1000) for 20 min. Localizations of LPG was determined as previously described (19). To label acidocalcisomes, stationary day 3 promastigotes were permeabilized with ice-cold ethanol and stained with rabbit anti-*T. brucei* VP1 antiserum (1:800) (42) for 30 min, followed by goat anti-rabbit IgG-Texas Red antibody (1:1000). DNA stain was performed with 1.5 μg/ml of Hoechst 33342 for 10 min. Images were acquired using an Olympus BX51 Upright Fluorescence Microscope equipped with a digital camera. To quantify the overlap between LSO-GFP and anti-BiP staining, 30 randomly selected cells were analyzed using Image J JACoP (Just Another Colocalization Plugin) (69). The fluorescence intensity of TbVP1 staining in WT, *lso*^*–*^ and *lso*^*–*^ /+*LSO* was measured from metacyclic-like cells (100 each) using Image J.

### Cell growth, stress response, and drug sensitivity

To measure parasite growth under the regular condition, log phase promastigotes were inoculated in M199 medium (pH 7.4) at 1.0 × 10^5^ cells/ml and incubated at 27 °C. Culture densities were determined daily using a hemocytometer. Percentage of round cells, dead cells, and metacyclics in stationary phase were determined as previously described (19). Parasite growth under the acidic condition was determined using an acidic M199 medium (same as the complete M199 medium except that the pH was adjusted to 5.0 with hydrochloric acid).

To assess cell viability under stress, mid log phase promastigotes were incubated in complete M199 medium (pH 7.4) at 37 (heat stress), in an acidic M199 medium (pH 5.0) at 27 (acidic stress) or in phosphate-buffered saline (PBS) at 27 (starvation stress). Cell viability over time was determined by flow cytometry after staining with 5 μg/ml of propidium iodide.

To determine sensitivity to drugs, log phase promastigotes were inoculated in complete M199 medium at 2.0 × 10^5^ cells/ml with different concentrations of Amp B (0.01-0.6 µM) or ITZ (0.01-13 µM). Percentages of growth were calculated after 48 hours by comparing culture densities from drug-treated cells to cells grown in absence of drugs (18).

To determine sensitivity to detergent, log phase promastigotes were inoculated in complete M199 medium with 0.0125% Triton X-100 at 2.0 × 10^5^ cells/ml. Cells viability was measured at different time points by flow cytometry after staining with propidium iodide.

### Intracellular pH measurement

Intracellular pH was measured using a pH sensitive fluorescent indicator, BCECF-AM (70). Briefly, promastigotes were inoculated in the regular (pH 7.4) medium or acidic (pH 5.0) medium at 1.0 × 10^5^ cells/ml as described above. After 3 days, 1.0 × 10^7^ cells were washed once with PBS and resuspended in 500 μl of buffer A (136 mM NaCl, 2.68 mM KCl, 0.8 mM MgSO_4_, 11.1 mM glucose, 1.47 mM KH_2_PO_4_, 8.46 mM Na_2_HPO_4_, 1 mM CaCl_2_, and 20 mM HEPES, pH 7.4) with 10 µM of BCECF-AM. After 30 min incubation, cells were washed twice and resuspended in buffer A. The emission intensity at 535 nm was measured using a microplate reader when samples were excited at 490 nm and 440 nm at the same time. The fluorescence intensity ratio (emission intensity at 535 nm when excited at 490 nm/emission intensity at 535 nm when excited at 440 nm) was converted into intracellular pH value using a calibration curve, which was generated by measuring fluorescence intensity ratios of cells prepared in pH 5.0-pH 8.0 of buffer A containing 10 μM of BCECF-AM and 5 μg/ml of the ionophore nigericin (71, 72).

### Acidocalcisome isolation and analysis of short chain and long chain polyphosphate

Acidocalcisome fractions were isolated from log phase and stationary phase promastigotes as described for *T. brucei* and *T. cruzi* (73). The amounts of short chain and long chain polyphosphate in acidocalcisome fractions were determined as previously described (74).

### Western blots

To determine LPG and GP63 expression, promastigotes were washed once in PBS and resuspended at 5.0 × 10^7^ cells/ml in 1 x SDS sample buffer. Samples were boiled for 5 min and resolved by SDS-PAGE, followed by immunoblotting with mouse-anti-*L. major* LPG monoclonal antibody WIC79.3 (1:1000) (75), mouse-anti-GP63 monoclonal antibody #235 (1:1000) (76), mouse-anti-*L. donovani* LPG monoclonal antibody CA7AE (1:500) or mouse-anti-α-tubulin antibody (1:1000), followed by a goat anti-mouse IgG-HRP (1:2000). To examine the expression of LSO-GFP, immunoblotting was performed using a rabbit anti-GFP antiserum (1:1000) followed by a goat anti-rabbit IgG-HRP (1:2000). To confirm LPG purification, 5 μg of purified LPG isolated from each strain was subjected to immunoblotting as described above using monoclonal antibody WIC79.3 (1:1000).

### Glycoconjugate extraction, purification, preparation and fluorophore-assisted carbohydrate electrophoresis (FACE)

LPG and GIPLs from *Leishmania* promastigotes (2-4 x 10^10^ cells each) were extracted in solvent E (H_2_O/ethanol/diethylether/pyridine/NH_4_OH; 15:15:5:1:0.017) and chloroform: methanol:water (10:10:3), respectively. The extracts were dried by N_2_ evaporation, resuspended in 0.1 M acetic acid/0.1 M NaCl, and applied to a column of phenyl-Sepharose (2 mL), equilibrated in the same buffer. LPG and GIPLs were eluted using solvent E (77).

To prepare LPG repeat units, the LPG samples were depolymerized by mild acid hydrolysis (0.02 M HCl, 5 min, 100 □). This would generate a mixture of phosphorylated repeat units and core-PI anchor which were separated after n-butanol:water (2:1) partitioning. Repeat units were collected in the aqueous phase and dephosphorylated with alkaline phosphatase in 15 mM Tris–HCl, pH 9.0 (1 U/ml, 16 h, 37 □). After enzymatic treatment, the repeat units were desalted by passage through a two-layered column of AG50W-X12 (H^+^) over AG1-X8 (acetate) (53). GIPLs were depolymerized after strong acid hydrolysis (2 M trifluoroacetic acid, 3 h, 100 □) to obtain neutral monosaccharides and samples were dried in a speed-vac and acid was removed by toluene wash (twice) under N_2_. GIPL samples were resuspended in 500 mL of water and desalted as described above (57).

LPG repeat units and GIPL monosaccharides were then subject to fluorophore-assisted carbohydrate electrophoresis (FACE) analysis. Purified LPG samples were fluorescently labeled with 8-aminonaphthalene-1,3,6-trisulfate and subjected to FACE analysis and the gel was visualized by an UV imager as described (53). To determine the monosaccharide composition of GIPLs, depolymerized and desalted GIPL samples were fluorescently labeled with 0.1 M 2-aminoacridone in 5% acetic acid and 1 M cyanoborohydride. Labeled sugars were subjected to FACE and the gel was visualized under UV light. Oligo-glucose ladders (G1–G7) and monosaccharides (D-galactose, D-glucose and D-mannose) were used as standards for oligosaccharides and monosaccharide gels, respectively (57).

### Mitochondria membrane potential, mitochondrial ROS and oxygen consumption

Mitochondria membrane potential was determined as described previously (18). Log or stationary phase promastigotes were resuspended at 1.0 × 10^6^ /ml in PBS with 100 nM of TMRE. After incubation at 27 °C for 15 min, cells were washed once with PBS and analyzed by an Attune Acoustic flow cytometer. To evaluate superoxide accumulation in mitochondria, promastigotes were resuspended in PBS at 1.0 × 10^7^ /ml and labeled with 5 µM of MitoSox for 25 min. Cells were washed once with PBS and analyzed by flow cytometry (18). To determine the oxygen consumption rate, log phase promastigotes were resuspended in a respiration buffer (Hank’s Balanced Salt Solution with 5.5 mM of sodium pyruvate, 5.5 mM of 2-deoxy-D-glucose) at 2.0 × 10^7^ /ml and oxygen consumption was measured with 1 μM of MitoXpress as described (63). WT parasites treated with 10 μM of antimycin A were included a negative control (63).

### Mouse footpad infection

BALB/c mice (female, 7–8 weeks old) were purchased from Charles River Laboratories International (Wilmington, MA). Mice were housed and cared for in a facility operated by the Animal Care and Resources Center at Texas Tech University. Procedures involving live mice were approved by the Animal Care and Use Committee at Texas Tech University (PHS Approved Animal Welfare Assurance No. A3629-01). To determine parasite virulence, 2.0 × 10^5^ metacyclics or 2.0 × 10^4^ lesion-derived amastigotes were injected into the footpad of each mouse (5 mice per group). Progression of footpad lesions were monitored weekly using a Vernier caliper. Anesthesia was applied via isoflurane inhalation during footpad injection and measurement. Euthanasia was achieved by CO_2_ asphyxiation. Parasites loads from infected footpads were determined by the limiting dilution assay (78).

## Statistical analysis

All experiments were repeated at least three times except for the Southern blot. All the graphs were made using Sigmaplot 13.0 (Systat Software Inc, San Jose, CA). Differences between two groups were determined by the Student’s *t* test. *P* values indicating statistical significance were grouped into values of <0.05 (*), <0.01 (**) and <0.001 (***).

## ACKNOWLEDGMENTS

We thank Dr. W. Robert McMaster (University of British Columbia, Vancouver, Canada) and Dr. Jay Bangs (University at Buffalo, State University of New York, USA) for kindly providing the anti-GP63 monoclonal antibody #235 and the rabbit anti-*T. brucei* BiP polyclonal antiserum, respectively. We also thank Dr. Catherine Wakeman (Texas Tech University, Lubbock, Texas, USA) for usage on the BioTek synergy 4 fluorescence microplate reader. This work is supported by National Institutes of Health grants AI099380 (KZ), P30DK020579, P30DK056341, and P41GM103422 (Mass Spectrometry Resource of Washington University). The funders had no role in study design, data collection and interpretation, or the decision to submit the work for publication.

## SUPPLEMENTAL FIGURE LEGENDS

**Figure S1. Sterol biosynthesis pathway in *L. major* WT and *lso*^*-*^ parasites**. I-VII represent sterol intermediates or final products (underlined). I: lanosterol, II: zymosterol, III: fecosterol, IV: episterol, V: 5-dehydroepisterol, VI: ergosterol, VII: ergosta-7,22-dienol. C14DM: Sterol C14α demethylase. SMT: Sterol C24-methyl transferase. LSO: lathosterol oxidase.

**Figure S2. LSO-GFP expression complements *lso***^***−***^ **mutants**. (**A**) Whole cell lysates from log phase promastigotes of *lso*^*−*^*/+LSO-GFP* (clones #1 and #2), *c14dm*^*−*^*/+C14DM-GFP* and WT were analyzed by Western blot (upper panel: anti-GFP; lower panel: anti-α-tubulin). (**B**)-(**E**) Partial GC-MS spectra of lipids from WT (**B**), *lso*^*−*^ (**C**), *lso*^*−*^*/+LSO* (**D**) and *lso*^*−*^*/+LSO-GFP* (**E**) promastigotes. The blue dashed lines indicate the shift of peaks in *lso*^*−*^ mutants in comparison to WT and add-backs.

**Figure S3. Short chain and long chain phosphate contents are not changed in** *lso*^*−*^ **mutants**. Acidocalcisomes from log phase or day 3 stationary phase promastigotes of WT, *lso*^*−*^ and *lso*^*−*^*/+LSO* were isolated and adjusted to the equivalence of 1.3 x 10^9^ cells/ml. The concentrations of short chain polyphosphate (**A**) and long chain polyphosphate (**B**) were determined as previously described. Error bars represent standard deviations from 3 repeats.

**Figure S4. *Lso***^***–***^ **mutants exhibit mild mitochondrial defects**. (**A-B**) Log phase and stationary phase (day 1-4) promastigotes were labeled with 100 nM of TMRE for 15 min for mitochondrial membrane potential (**A**) or 5 µM of MitoSox Red for 25 min for mitochondrial ROS level (**B**). Mean fluorescence intensities were determined by flow cytometry. (**C**) Log phase promastigotes were resuspended in a respiration buffer (HBSS + 5 mM 2-deoxyglucose + 5 mM sodium pyruvate) and oxygen consumption over time was measured after labeling with 1 μM of MitoXpress. Error bars represent standard deviations from 3 experiments (*: *p* <0.05, **: *p* <0.01).

