## supplementary data for "Lathosterol oxidase (sterol C5-desaturase) deletion confers resistance to amphotericin B and sensitivity to acidic stress in *Leishmania major*"

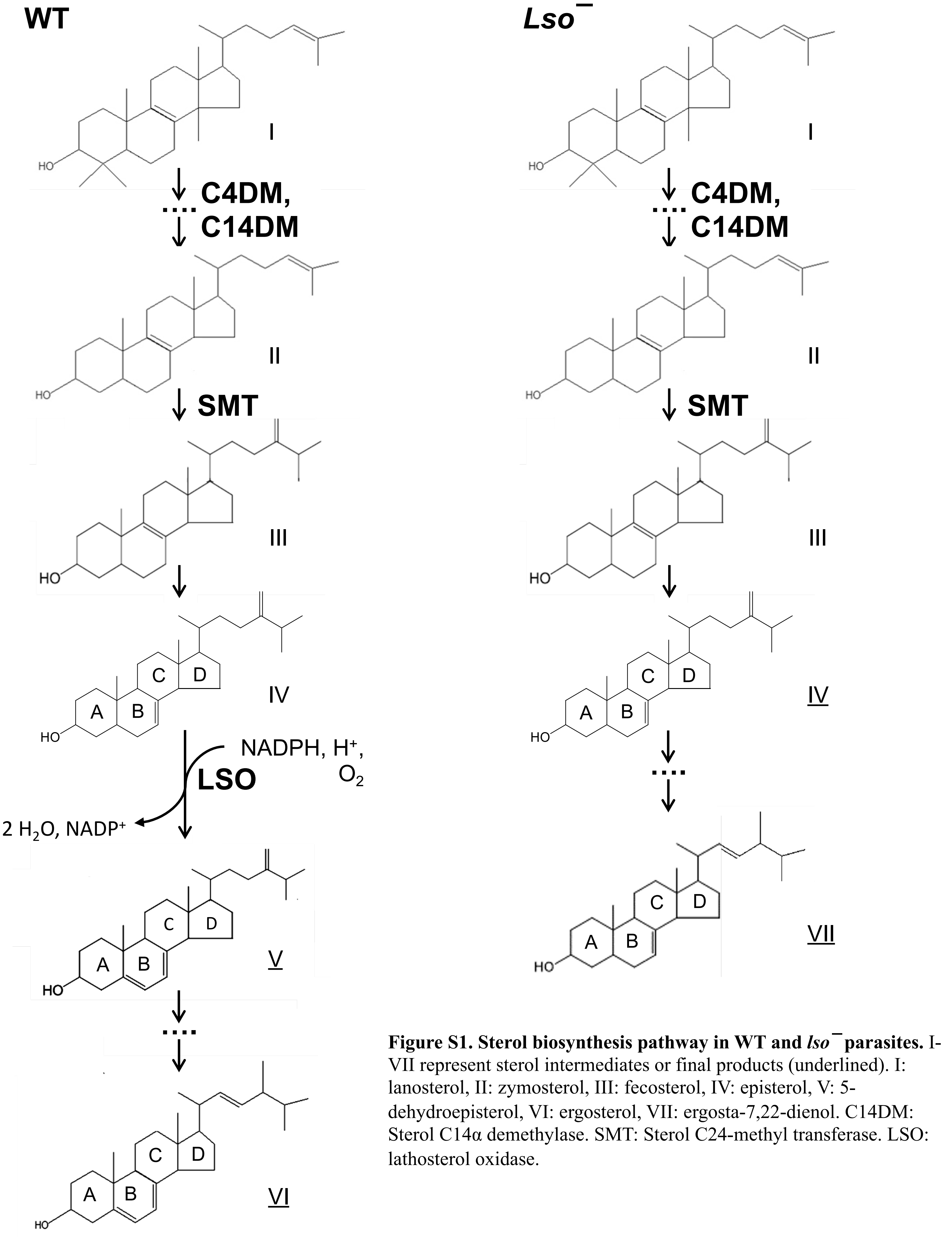


**Figure S1. Sterol biosynthesis pathway in WT and *lso*$^{-}$ parasites.** I-VII represent sterol intermediates or final products (underlined). I: lanosterol, II: zymosterol, III: fecosterol, IV: episterol, V: 5-dehydroepisterol, VI: ergosterol, VII: ergosta-7,22-dienol. C14DM: Sterol C14α demethylase. SMT: Sterol C24-methyl transferase. LSO: lathosterol oxidase.

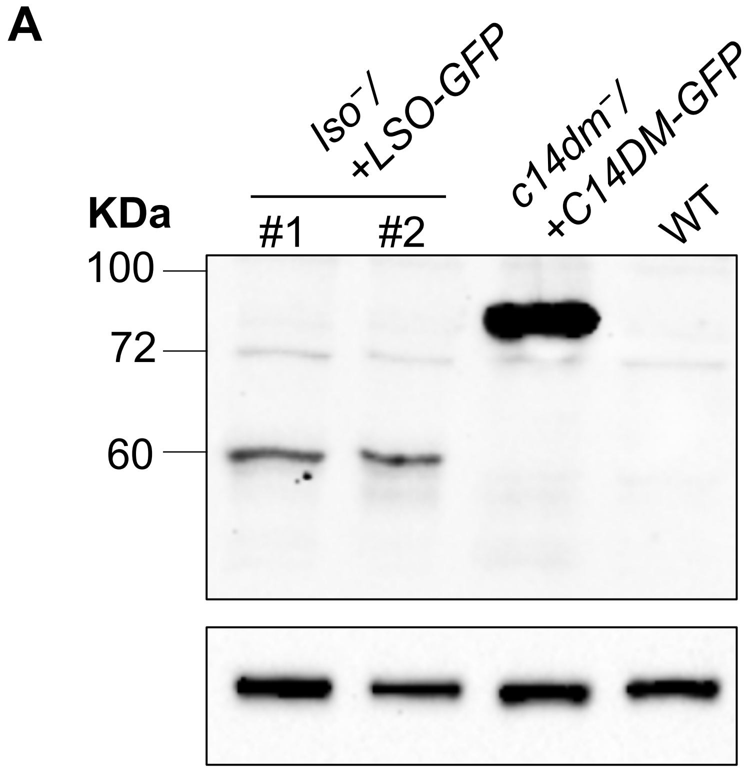


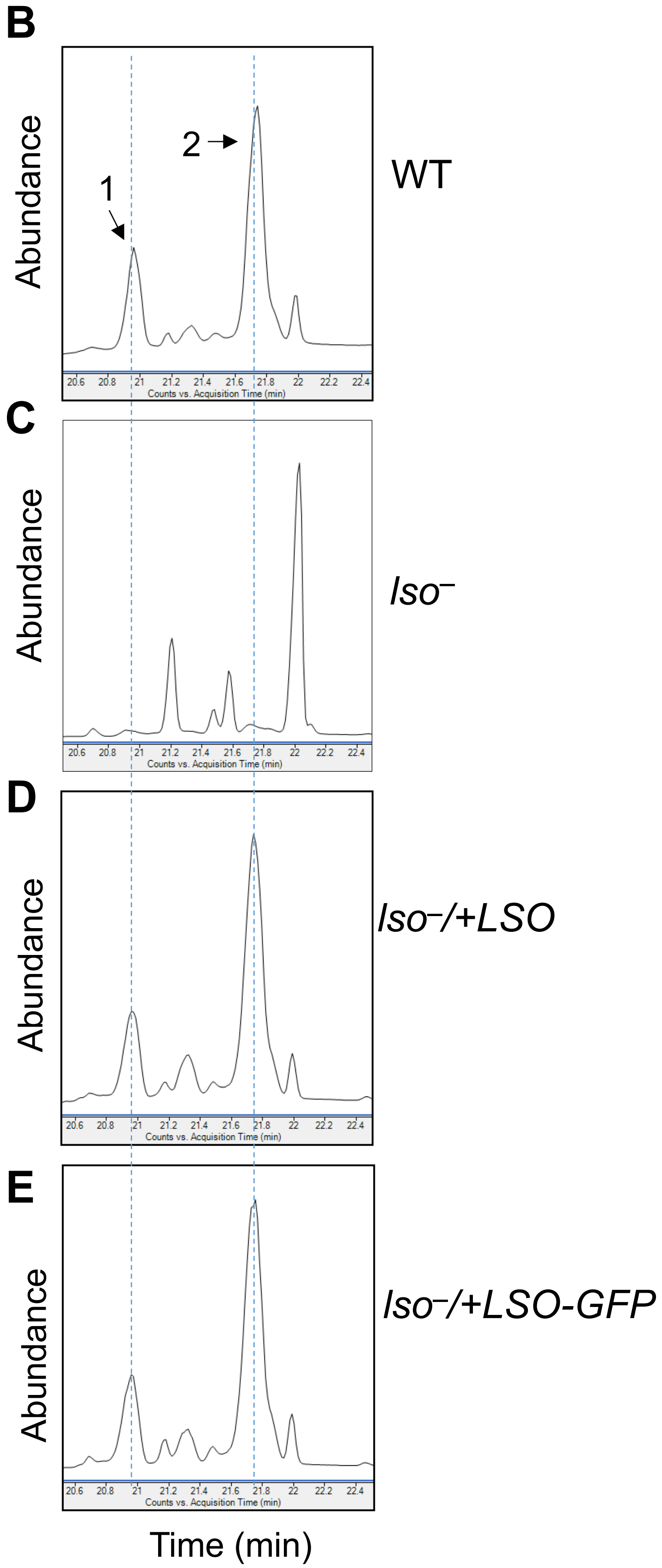


**Figure S2. LSO-GFP expression complements *lso*⁻ mutants.** (**A**) Whole cell lysates from log phase promastigotes of *lso⁻/+LSO-GFP* (clones #1 and #2), *c14dm⁻/+C14DM-GFP* and WT were analyzed by Western blot (upper panel: anti-GFP; lower panel: anti-α-tubulin). (**B**)-(**E**) Partial GC-MS spectra of lipids from WT (**B**), *lso⁻* (**C**), *lso⁻/+LSO* (**D**) and *lso⁻/+LSO-GFP* (**E**) promastigotes. The blue dashed lines indicate the shift of peaks in *lso⁻* mutants in comparison to WT and add-backs.

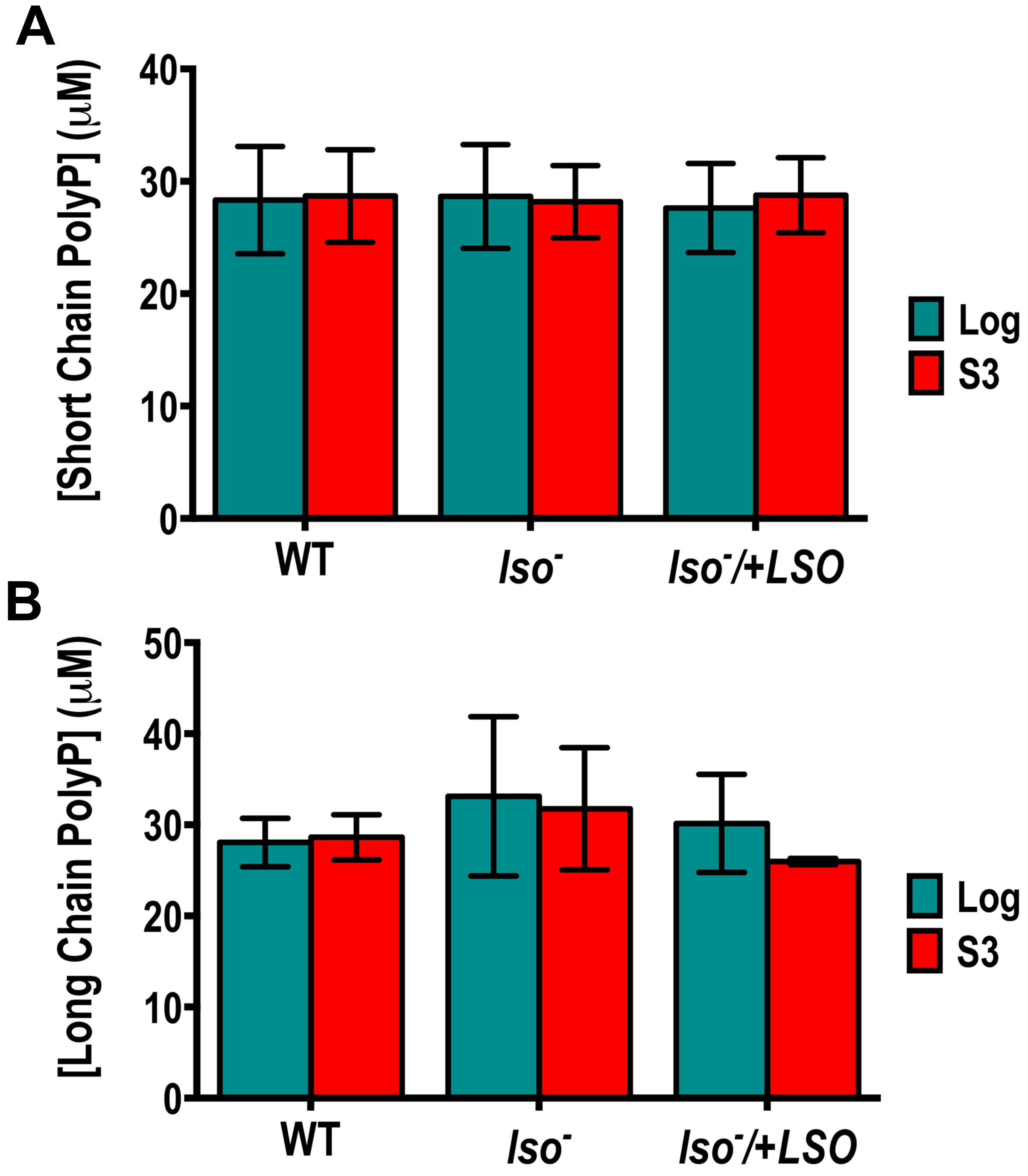


**Figure S3. Short chain and long chain phosphate contents are not changed in *lso*⁻ mutants.** Acidocalcisomes from log phase or day 3 stationary phase promastigotes of WT, *lso*⁻ and *lso*⁻*/+LSO* were isolated and adjusted to the equivalence of 1.3 x $10^9$ cells/ml. The concentrations of short chain polyphosphate (**A**) and long chain polyphosphate (**B**) were determined as previously described. Error bars represent standard deviations from three repeats.

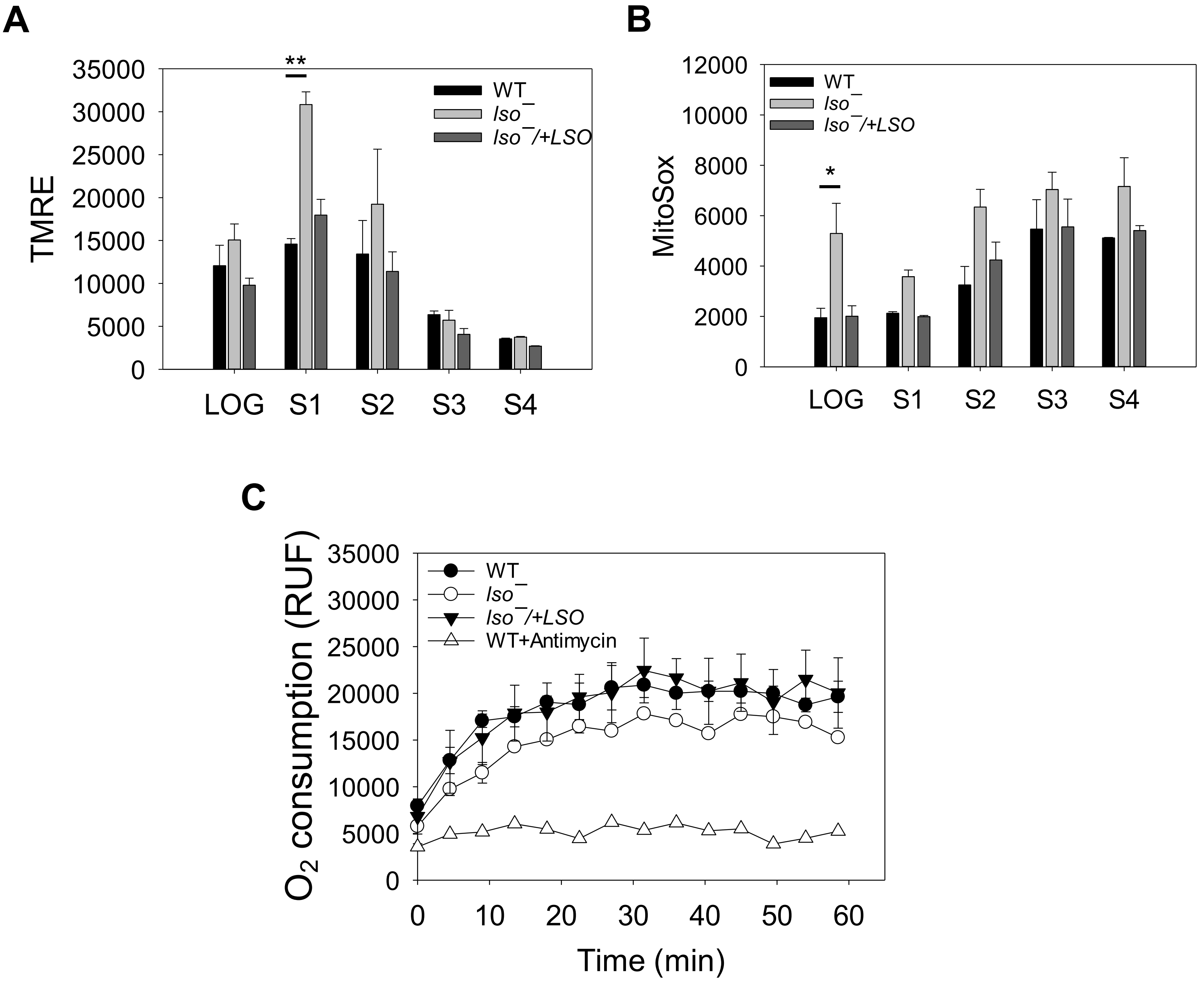


**Figure S4. *Lso*$^{-}$ mutants exhibit mild mitochondrial defects.** (**A-B**) Log phase and stationary phase (day 1-4) promastigotes were labeled with 100 nM of TMRE for 15 min for mitochondrial membrane potential (**A**) or 5 µM of MitoSox Red for 25 min for mitochondrial ROS level (**B**). Mean fluorescence intensities were determined by flow cytometry. (**C**) Log phase promastigotes were resuspended in a respiration buffer (HBSS + 5 mM 2-deoxyglucose + 5 mM sodium pyruvate) and oxygen consumption over time was measured after labeling with 1 µM of MitoXpress. Error bars represent standard deviations from 3 experiments (*: $p$ <0.05, **: $p$ <0.01).

**Table S1. List of oligonucleotides used in this study.**

| Primer # | Name | Purpose | Sequence |
|---|---|---|---|
| 645 | LSO-5'UTR-F-EcoRI | To amplify the 5’-flanking sequence | CATGATgaattcCACTGTACGTCGGCTCGTG |
| 646 | LSO-5'UTR-R-SpeI | To amplify the 5’-flanking sequence | GTCGCTactagtGCTTTCGAATGAGCCGGTG |
| 647 | LSO-3'UTR-F-SpeI-BglII | To amplify the 3’-flanking sequence | CGGACGactagtGGCTAGagatctGAGAAGGATCGGTCACCTTATTG |
| 659 | LSO-3'UTR-rev (HindIII) | To amplify the 3’-flanking sequence | GGCAGCaagcttCCTACGTACTTACTCGCAC |
| 649 | LSO-ORF-F-BamHI | To amplify the *LSO* ORF | TCAGTAggatccACCATGGACTTCGCCTTTCGCC |
| 650 | LSO-ORF-R-BamHI | To amplify the *LSO* ORF | GCACTggatccTTATGCGCCGACCTTCTTTGAGC |
| 651 | LSO-GFP-R-EcoRV | To generate the *LSO-GFP* fusion | GATGCTgatatcTGCGCCGACCTTCTTTGAGC |
| 653 | LSO-Probe 1-F | To generate the LSO probe for Southern blot | GAGTAACCGAGTCGGTTGTC |
| 654 | LSO-Probe 1-R | To generate the LSO probe for Southern blot | GTGAGAAGTGTCTCGACG |
| 655 | LSO-Probe 2-F | To generate the LSO probe for Southern blot | CTCTTCGCCTCCATCTCCTAC |
| 660 | LSO-Probe 2-R | To generate the LSO probe for Southern blot | GATCGCCAGAAGTTCACCAC |
| 848 | SCGs-F | For qRT-PCR analysis of SCGs | GAGGAGAACAATGTGCCAG |
| 849 | SCGs-R | For qRT-PCR analysis of SCGs | CCGCAGTGCCTCTGAGAG |
| 709 | 28S rRNA-F | For qRT-PCR analysis of 28S rRNA | AAGATGGACCGGCCTCTAGT |
| 710 | 28S rRNA-R | For qRT-PCR analysis of 28S rRNA | ATCCTTCCCCGCTCCAGTAT |

Sequences in lowercase represent restriction enzyme recognition sites.
